# Genetic encoding of climate-responsive stomatal developmental plasticity in tomato

**DOI:** 10.64898/2026.03.27.714625

**Authors:** Ido Nir, Alanta Budrys, Daniel Suraev, Hermann Prodjinoto, Joel Erberich, Jonathan Tirnover, Ella Zafrir, Yaarit Kutsher, N. Katherine Smoot, Dominique C. Bergmann

## Abstract

Flexible developmental programs enable plants to customize their organ size and cellular composition. In leaves of eudicots, the stomatal lineage produces two essential cell types, stomata and pavement cells, and plants can adjust the total numbers and ratios of these cell types in response to external cues. Central to this flexibility is the stomatal lineage-initiating transcription factor, SPEECHLESS (SPCH). Here we explore the mechanisms underlying SPCH’s involvement in environmental response. Using multiplexed CRISPR/Cas9 editing of *SlSPCH cis-*regulatory sequences in tomato, we identified variants with altered stomatal development responses to drought, light and temperature cues. By creating and live-cell tracking translational reporters of SlSPCH and its paralogues SlMUTE and SlFAMA, we revealed the corresponding cellular events that lead to the environmental change-driven responses in stomatal production and leaf form. Plants bearing the novel reporters and *SlSPCH* variants are powerful resources for fundamental and applied studies of tomato resilience in response to climate change.

## INTRODUCTION

In response to changing climates, plants adjust their physiology and growth. A conserved and well-studied response is the change in activity and production of stomata upon environmental perturbation. Stomata are cellular valves in the epidermal surfaces of aerial organs; a pair of stomatal guard cells (GCs) flank a pore whose aperture they modulate through turgor-driven cell swelling. Stomatal opening and closing responses occur within minutes of a change in light, water availability, carbon dioxide concentration [CO_2_], or temperature, and many of the signal transduction pathways for perception and response have been described in detail (reviewed in (1)). Over longer timescales, stomatal responses to environmental change are detected as changes in stomatal density (SD, stomata/unit area) or stomata index (SI, stomata/total epidermal cell number). Because stomata are preserved in fossils and herbaria, these developmental responses have been used as proxies for the composition of past climates (2, 3). Changes to SI and SD can also occur during the development of individual leaves of plants subjected to changes in light (4, 5) temperature or [CO_2_] (6–8) (Figure 1a).

**Figure 1.**
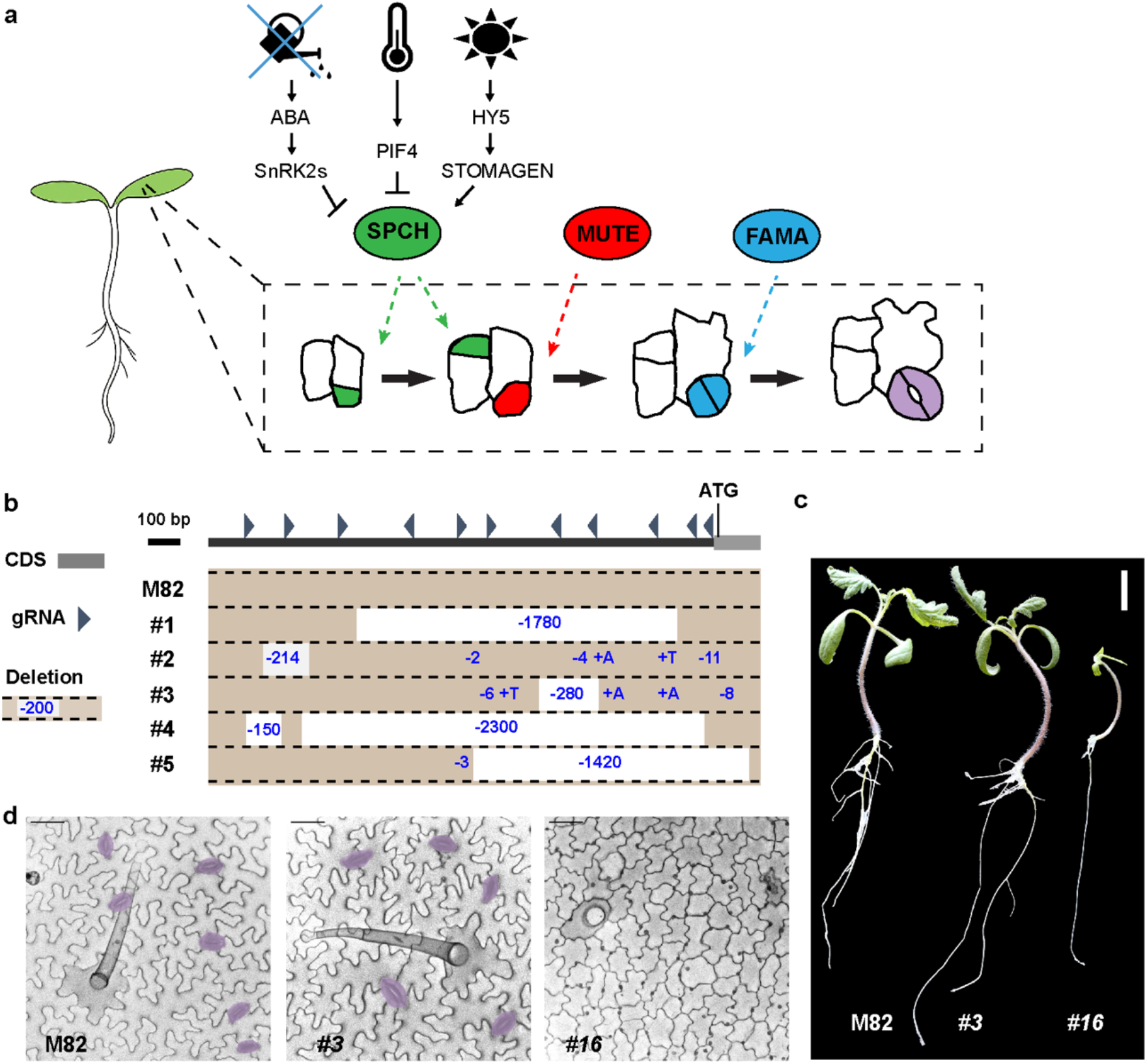
Generation of *SlSPCH* cis-regulatory mutations to identify inputs into flexible early stages. a) Cartoon of generalized dicot stomatal lineage including core transcription factors (ovals) color-coded by cell-type they are expressed in and regulate. Inside dashed box are representations of divisions within a clone of leaf epidermal cells. Green indicates an asymmetric stomatal lineage-initiating division. Red indicates a cell that has committed to stomatal fate, and blue and purple represent young and mature stomatal guard cells. SPCH and the green divisions are affected by environmental information relayed via transcriptional regulation of SPCH and/or via intracellular signaling cascades that phosphoregulate SPCH protein. b) Diagram of *SlSPCH cis*-regulatory mutagenesis scheme, with position of gRNAs 5’ of the SlSPCH start site and resultant deletion alleles characterized in detail in this paper (c) Seedling morphology and (d) cotyledon epidermis phenotypes in wildtype M82, *SlSPCH* cis-regulatory deletion line #3 and *SlSPCH* coding region deletion line #16. Stomata are false-colored purple in (d). Scale bars in (c) 20 mm, and in (d) 50 µm.

Genes responsible for stomatal production and pattern have been identified, typically first in *Arabidopsis thaliana*, with subsequent studies showing that a core set of transcription factors, receptors and signaling peptides are broadly used among plants (e.g. (9–16)). Among angiosperms, the plants that comprise the majority of natural and cultivated ecosystems, the transcription factors (TFs) SPEECHLESS (SPCH), MUTE, FAMA and their heterodimer partners ICE1/SCREAM and SCRM2 play key roles in the specification of stomatal precursors and in the differentiation of the stomatal guard cells and subsidiary cells (reviewed in (17, 18)).

Temperature, light, atmospheric [CO_2_] and water availability affect stomatal function and production, with increasing light and water typically leading to an increase in SD and SI (19). Due to its role regulating the initiation and proliferation of stomatal precursors, SPCH is often the ultimate target downstream of environmental cues (Figure 1a). For example, direct regulation of *SPCH* transcription in response to warm temperature is mediated by PHYTOCHROME INTERACTING FACTOR4 (PIF4) (6) and *SPCH* transcript levels and/or SPCH protein stability are impacted by changes in light intensity (20–23). Stomatal development is also regulated systemically by plant hormones with the drought hormone abscisic acid (ABA) having a major impact. ABA can negatively regulate stomatal production via the degradation of SPCH, leading to reduced stomatal number and reduced water loss (23). Although *SPCH* responds to many internal and environmental signals, little is known about the specific regulatory sites in its *cis*-regulatory regions that respond to the different cues.

Advances in precision genome editing have enabled new ways of modulating gene activities, with the potential to also reveal endogenous regulatory mechanisms. Multiplex CRISPR/Cas9 mediated editing of tomato gene *cis*-regulatory regions, for example, revealed enhancers responsible for architecture and fruit traits that mimic those bred into specialized commercial tomato lines, as well as generating novel fruit morphologies (24). Given these technical innovations and the placement of *SPCH* in stomatal and leaf development networks, *SPCH* emerged as an ideal candidate for *cis*-regulatory dissection aimed at uncovering elements that confer responsiveness to environmental inputs. Here we show that *SlSPCH* cis-regulatory alleles exhibit unique and differential stomatal “set points”, as well as divergent responses to environmental change. Another lesson from recent engineering and comparative studies in angiosperms, however, is that developmental regulators and pathways shift (e.g. (25)) and it is crucial to assess edited gene function in the context of species-specific programs. Therefore, to create a platform to examine stomatal plasticity in response to environmental change in tomato, we combined the *SlSPCH cis*-element dissection with complementary live-cell imaging approaches. We generated reporters to track cell lineages track and to follow expression of SlSPCH, SlMUTE and SlFAMA in the tomato leaf epidermis over time, thereby defining the precise cellular pathways tuned by environmental inputs. Together these new data and tools reveal genetic and cellular mechanisms by which tomato adjusts stomatal production in response to environmental cues.

## RESULTS

### Mutagenesis of the *SlSPCH* cis-regulatory region generates small deletions that display a range of stomatal phenotypes

Because SPCH plays a central role in stomatal lineage initiation and flexibility in other species, we targeted *SlSPCH* for mutagenesis of its *cis*-regulatory region, hypothesizing that we could identify elements that rendered its expression tunable to environmental cues. We expected that the complete loss of SlSPCH activity would eliminate stomata and be lethal, thus we employed a CRISPR/Cas9-based multiplex strategy (24) to make a series of small deletions in the 5’ regulatory region of *SlSPCH*. The nearest protein coding gene lies more than 7.5 Kb upstream of the *SlSPCH* translational start site (Figure S1a), but ATAC-seq profiles of open chromatin suggested that for most tomato genes, the region ∼3Kb upstream of the start site is likely to contain key regulatory regions (26), and indeed in this smaller region we find numerous predicted target sites for transcription factors with roles in developmental, environmental and hormone-responsive regulation (Figure S1a). We generated 11 sgRNAs, as evenly distributed as sequence would allow, and generated a series of *SlSPCH* cis-regulatory alleles that range from 2bp to ∼2.5Kbp (Figure 1b and Table S1), as assayed by genotyping of T1 plants. Plants with deletions spanning more than ∼100 bp were selected and self-pollinated to obtain homozygous lines for all subsequent experiments.

From the initial mutagenesis, we identified 15 independent lines with distinct *cis*-regulatory mutations (Table S1). Because CRISPR/Cas9 mutagenesis can lead to a spectrum of mutations, we also recovered plants where deletions disrupted the *SlSPCH* coding region (e.g., #16, Figure S1d-e). *SlSPCH* #16 was pale, lacked stomata and arrested as a seedling (Figure 1c-d), similar to *Arabidopsis spch* null mutants (27), confirming that *SlSPCH* is essential for stomatal lineage initiation. Our main interest, however, was conditional mutants that failed to adjust stomatal production in response to environmental cues, but whose overall size and leaf morphology was not substantially different from M82 under standard growth conditions. From these lines, we prioritized a set whose mutations tiled across the *SlSPCH* cis-regulatory region, ultimately choosing five for detailed study of developmental and physiological responses to altered water availability, light and temperature conditions (Figure 1b and Table S1).

### SlSPCH *cis*-regulatory mutants display altered responsiveness to drought

Drought resilience is historically linked to the ability of plants to alter stomatal aperture in response to ABA (28). ABA, however, also affects stomatal development (29) and ABA initiates changes in gene expression through a set of conserved transcription factors that bind ABRE motifs (ACGTGG/TC), DRE/CRT dehydration-responsive elements (A/GCCGAC) (30) and/or MYB-type drought-associated (MBS) motifs (C/TAACNA/G) (31) in the *cis*-regulatory regions of target genes. In the full-length *SlSPCH cis*-regulatory region, coexistence of ABRE, DRE/CRT, and MYB-type elements (Figure S1a) suggests that *SlSPCH* expression may integrate information from several drought signaling pathways, mirroring the regulatory architecture of known stress-responsive plant promoters. Several of our deletions disrupt or rearrange these ABA and/or drought associated motifs (Figure S1b), enabling us to test the hypothesis that plants reduc e stomatal numbers under long-term drought via these known pathways.

We assessed the impact of *SlSPCH* cis-regulatory mutation on whole plant transpiration and on SI in response to water deficit. Briefly, M82 controls and the *SlSPCH* cis-regulatory mutants were grown under controlled conditions of sufficient water or water deficiency of 30% relative soil water content (RSWC) for three weeks. All plants were then grown under a sufficient water regime for a one-week recovery period (Figure 2a, and methods). We measured the daily whole plant transpiration rate and the SI of expanded 5^th^ leaves in the four-week-old plants. In M82 controls, we observed a significant decrease in SI (Figure 2b) and daily transpiration (Figure 2c) in plants grown in 30% RSWC compared to well-watered plants. The *SlSPCH* cis-regulatory variants showed diverse responses in SI and daily transpiration. Line *#1* which deletes a proximal ABRE and three MBS elements, had a slightly lower SI than M82 under all conditions, but was similar to M82 in magnitude and direction of SI and daily transpiration responses to 30% RSWC (Figure 2b-c). Line *#2*, which deletes a distal ABRE and MBS element, line *#4* which deletes a proximal ABRE and three MBS, but creates a new ABRE, and line *#3* which deletes two proximal MBS elements, failed to significantly reduce transpiration in response to 30% RSWC (Figure 2c) indicating that these lines were insensitive to long-term drought. The fact that their SI was also insensitive (Figure 2b) suggests that at least part of this drought response is attributable to mis-regulation of stomatal production, though our data suggest that ABRE and MBS elements are not crucial for this mechanism (Figure S1b). Line *#5* had a significantly lower SI than M82 and the other *SlSPCH* mutants under both drought and well-watered conditions (Figure 2b). Daily transpiration rates in line *#5* were also insensitive to drought (Figure 2c), but interestingly, these rates were similar to that of lines with higher SI suggesting compensation through altered stomatal activity (32).

**Figure 2.**
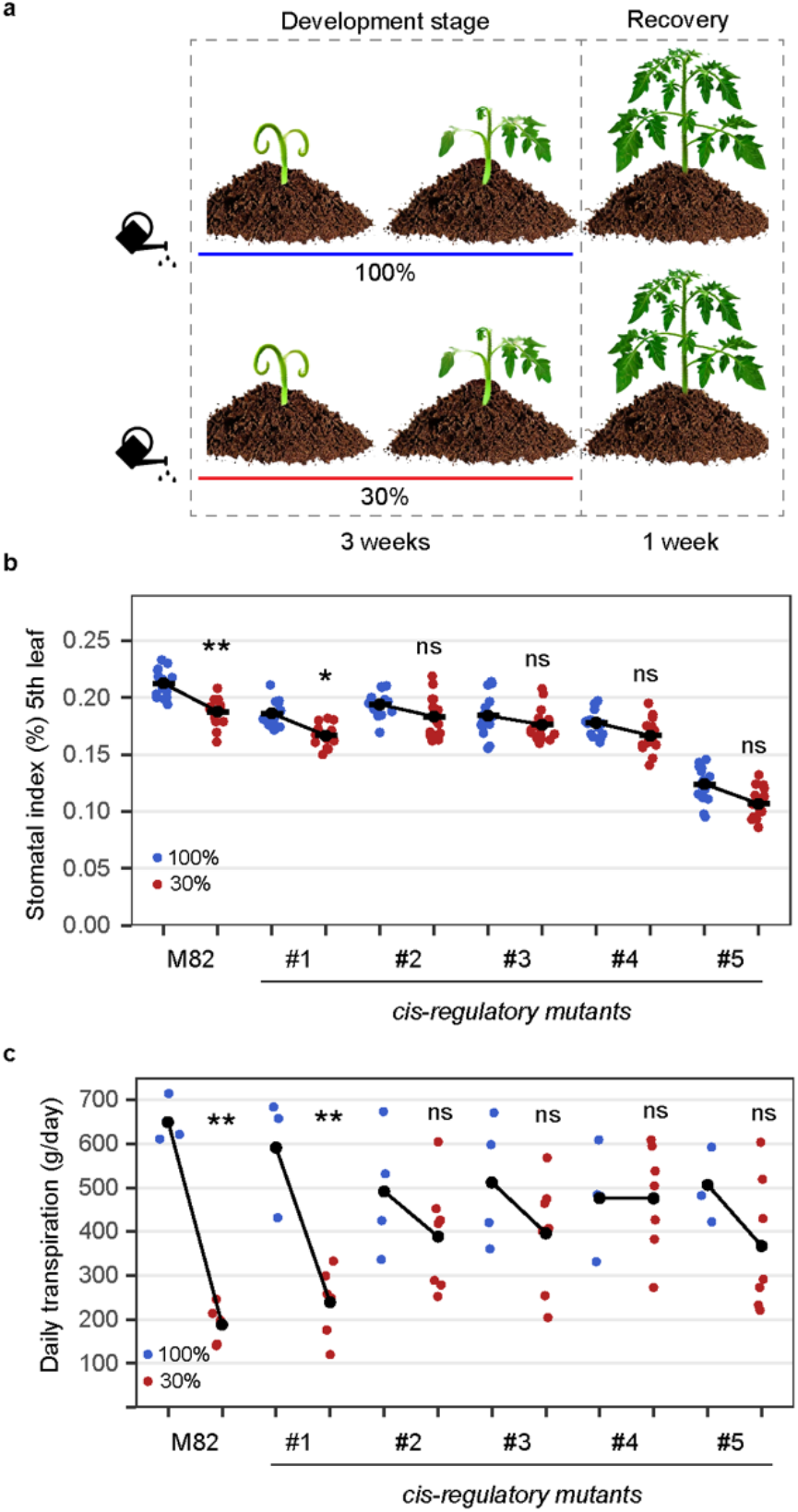
Stomatal response to changes in water availability is altered in some *SlSPCH* cis-regulatory mutants. (a) Diagram of the water deficit experiment, illustrating the 3-week development stage where soil grown plants generate leaves under either 100% or 30% irrigation levels, followed by a one-week recovery stage at 100% irrigation. (b Plot of stomatal index (SI) in 5^th^ leaf showing the developmental response to full irrigation (100%, blue) and water deficit (30%, brown) conditions in M82 and *SlSPCH cis*-regulatory mutants. (c) Plot of daily transpiration in mature plants showing the physiological capacity response to full irrigation (100%, blue) and water deficit (30%, brown) conditions in M82 and *SlSPCH cis*-regulatory mutants. Statistical tests in (b-c) are represented as mean ± 95% confidence interval. Bonferroni-corrected p values from Mann-Whitney U test are *P < 0.05; ***P < 0.001; ****P < 0.0001. n.s.: P > 0.05, not significant. Sample sizes in (b) are n = 8-16 0.4mm2 fields from 6-8 true leaves.

Considered together, these analyses reveal robust, allele-specific effects of *SlSPCH cis*-regulatory mutations on drought responses in stomatal development and at the whole-plant level. This endpoint analysis, however, does not capture the underlying cellular dynamics of stomatal lineage regulation. We therefore turned to a more careful analysis of mutant and reporter-bearing seedlings grown under controlled light and temperature conditions, enabling a high-resolution analysis of *SlSPCH-*dependent lineage behavior in response to these environmental cues.

### *SlSPCH* cis-regulatory mutants display altered responsiveness to changes in light intensity

As the main organs of photosynthesis, leaves have finely tuned responses to changes in light quality and intensity. Along with increasing their mesophyll and chloroplast production, most plants respond to elevated light with an increase in stomatal index (SI) and in total numbers of stomata per leaf. In Arabidopsis, light-mediated promotion of stomatal development is accompanied by higher levels of SPCH. The light responsive transcription factor ELONGATED HYPOCOTYL 5 (HY5) acting in the mesophyll initiates a pathway that ultimately results in higher SPCH protein levels and more stomata (4). HY5 is also expressed in the epidermis (33). It has been suggested, though not experimentally confirmed, that *SPCH* transcription would also be regulated by light (22) and there is a predicted HY5 binding site deleted in lines *#1 and #4* (Figure S1b).

To establish a baseline light response for tomato line M82 (with intact *SlSPCH*), we grew M82 plants at 26°C and two light intensities, ∼130 µmol-photons m^-2^ s^-1^ and ∼1300 µmol-photons m^-2^ s^-1^. We quantified SI, SD and leaf area (Figure 3a-b and S2a-b) as measures of developmental response. As expected, M82 plants increased their SI and SD in the higher light condition. We then tested our *SlSPCH* cis-regulatory deletion mutants in the same conditions and found a variety of responses (Figure 3a). Lines #1-#3 have similar SIs to M82 at low light but exhibit a dampened response to higher light intensity. Line #4 has a lower SI at low light, but is similar to M82 at high light, and therefore has an exaggerated response. Line #5 exhibits both a lower SI at low light and no response to increase in light intensity (Figure 3a). Similar trends are seen when SD is calculated (Figure 3b and S2b) except that line #5 does show an increase in SD. SD measurements are sensitive to changes in overall leaf size which typically changes when plants are grown in high light (Figure S2a) whereas SI, as a ratio of cell types is less so. Taken together these phenotypes suggest that alteration of the cis-regulatory region of *SlSPCH* can render stomatal production differentially sensitive to light, but that the HY5 binding site is not essential for this regulation.

**Figure 3.**
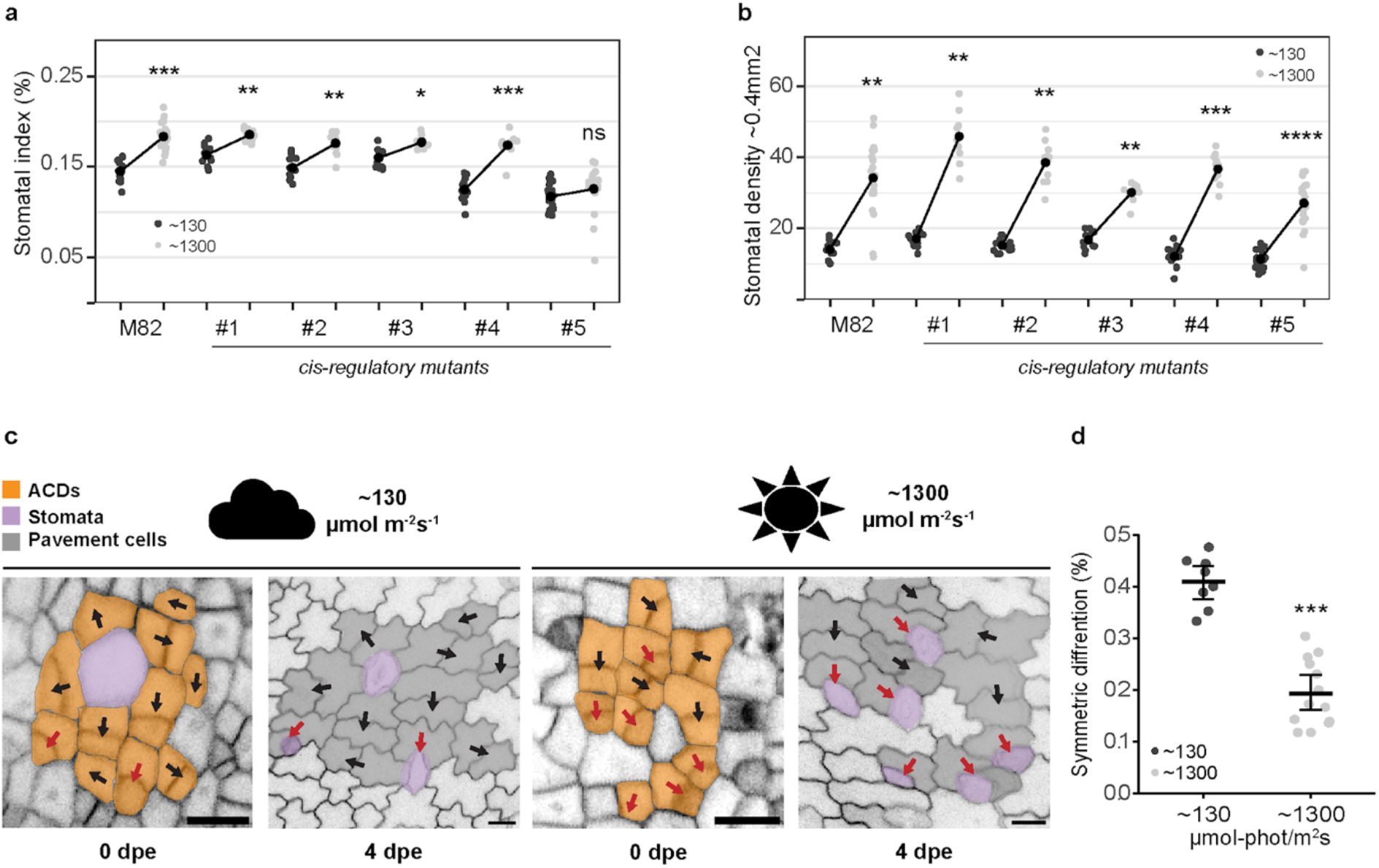
Stomatal response to changes in light intensity occurs via a lineage-exit strategy and is altered in some *SlSPCH* cis-regulatory mutants. (a-b) Plot of stomatal index (SI, a) and stomatal density (SD, b) response to low (black) and high (pale grey) light conditions in M82 and *SlSPCH cis*-regulatory mutants. (c) lineage tracing of asymmetrically dividing cells and their fate outcomes from confocal images of M82 cotyledons expressing epidermal plasma membrane reporter *ML1p:RCI2A-NeonGreen*. Red arrows mark ACDs that yield stomata (purple) and black arrows indicate ACDs that lead to two pavement cells (symmetric differentiation). Scale bars represent 20 µm. (d) Plot of shift in the number of ACDs that yield two pavement cells in low and high light. Statistical tests in (a), (b) and (d) are represented as mean ± 95% confidence interval. Bonferroni-corrected p values from Mann-Whitney U test are *P < 0.05; **P < 0.01; ***P < 0.001; ****P < 0.0001. n.s.: P > 0.05, not significant. Sample sizes in (a) and (b) are n = 8-21 0.4mm^2^ fields from 3-5 cotyledons. Sample sizes in (d) are n = 8-14 0.4mm^2^ fields from 3-4 cotyledons.

We next determined how SlSPCH-guided divisions in the stomatal lineage could change SI in response to light. In Arabidopsis, a shift in the relative frequency of amplifying to spacing asymmetric divisions leads to an increase in SI (34). However, in tomato, spacing divisions are infrequent (35), so it was unclear what cellular mechanisms were available for tomato stomatal lineages to alter SI. We therefore imaged the epidermis of abaxial cotyledons expressing the epidermal plasma membrane reporter ML1p::RCI2A-NeonGreen (35) every two days to trace asymmetrically dividing precursor cells to their final fate outcome (Figure 3c). From these time courses, we found that the major mechanism responsible for altering SI was a shift from asymmetric divisions that yield one stoma and one pavement cell to asymmetric divisions that produced two pavement cells (“symmetric differentiation”, Figure 3d). Altering stomatal ratios by altering the fate or differentiation outcome of early stomatal lineage divisions rather than changing the frequency of those divisions has been termed a “lineage exit” strategy (35); here we show that lineage exit is used to decrease SI in low light.

### SlSPCH cis-regulatory mutants display altered responsiveness to changes in temperature

Plants also respond to elevated temperature by changing stomatal behavior and production (7, 36). In Arabidopsis, *SPCH* transcription is repressed by the warm-temperature-induced PIF4 transcription factor (6). Adapting the temperature-shift protocols in (6) to account for the standard growth temperatures of tomato (26°C), we measured the M82 response to elevated (34°C) temperature. Although leaf size responses to growth at elevated temperature were similar between Arabidopsis and tomato (Figure S2c), SD and SI differed in the direction of response. As reported in (6), higher temperatures suppressed SI in Arabidopsis, whereas in tomato M82, they led to higher SI and SD (Figure 4a and S2d).

**Figure 4.**
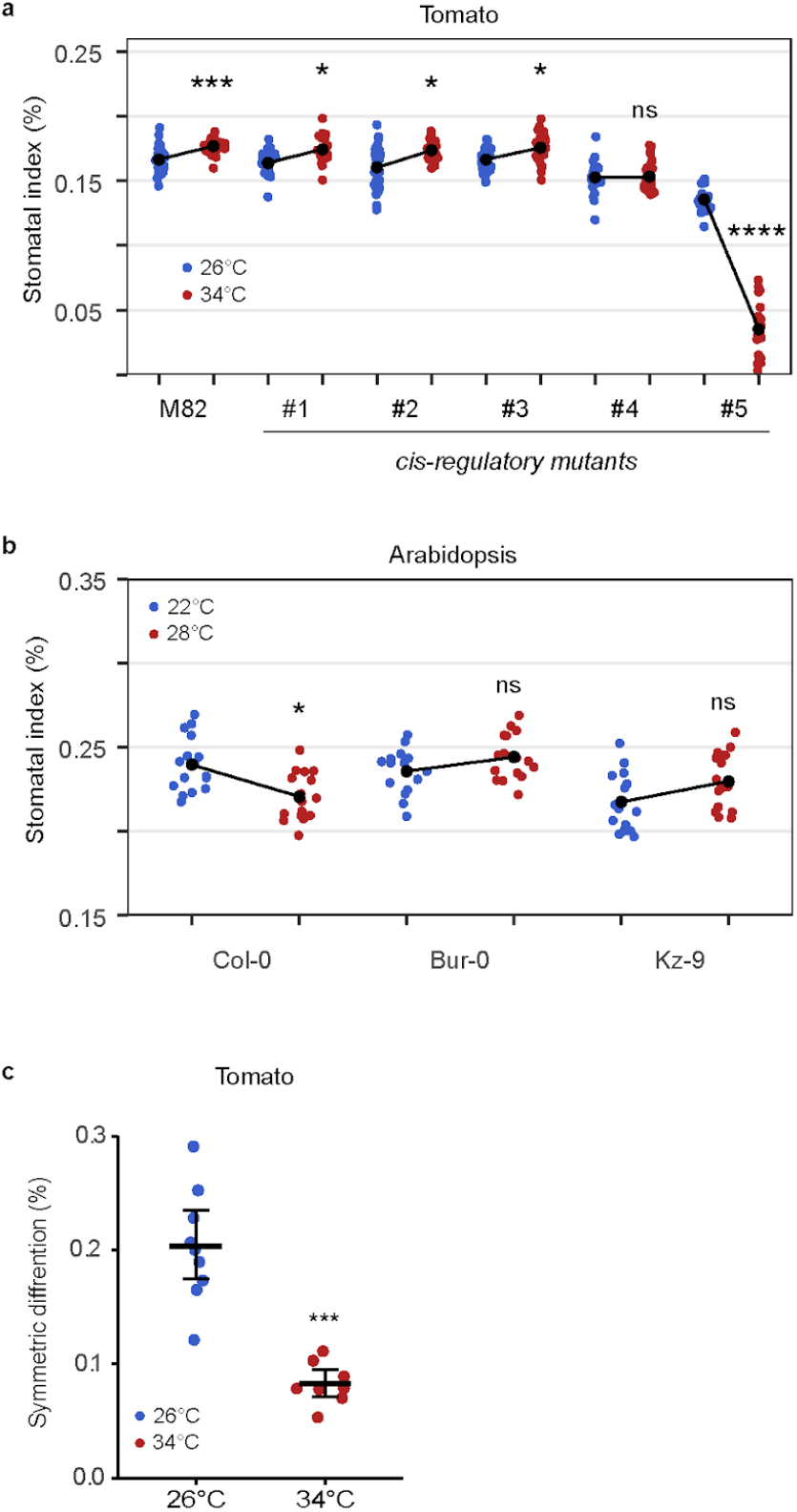
Stomatal response to changes in temperature occurs via a lineage-exit strategy and is altered in some *SlSPCH* cis-regulatory mutants and among Arabidopsis accessions. (a) Plot of SI of M82 and *SlSPCH cis*-regulatory mutants in response to growth at 26°C (blue) or 34°C (red) (b) Plot of SI of Arabidopsis accessions in response to growth at 22°C or 28°C. (c) Plot of shift in the number of tomato stomatal lineage ACD that produce two pavement cells instead of one stoma and one pavement cell (symmetric differentiation) in plants grown at 26°C and 34°C. Statistical tests in (a-c) are represented as mean ± 95% confidence interval. Bonferroni-corrected p values from Mann-Whitney U test are *P < 0.05; ***P < 0.001; ****P < 0.0001. n.s.: P > 0.05, not significant. Sample sizes in (a) n = 20-30 0.4mm^2^ fields from 4-6 cotyledons. Sample sizes in (b) n= 15-16 cotyledons each from an individual plant. Sample sizes in (c) are n = 8-9 fields 0.4mm^2^ from 4-6 cotyledons.

The opposite responses of tomato and Arabidopsis to increased temperature intrigued us. While there are numerous reports of different plants exhibiting stomatal opening and closing responses to changing temperature (37–39), stomatal production or pattern responses are less clear. The Arabidopsis accession used in (6) was Col-0, but there are Arabidopsis accessions from diverse climates and with diverse life history traits. We repeated the Arabidopsis temperature-shift experiments with Col-0 (Germany) and accessions Bur-0 (Ireland) and Kz-9 (Kazakhstan). As before, high temperature results in significantly lower SI in Col-0, but this reduction is not seen in Bur-0 or in Kz-9 (Figure 4b). Additional measurements of SD and leaf area indicate that there is a diversity of responses, and Col-0 alone is not sufficient to fully encompass the “Arabidopsis” response (Figure S2e-f).

Returning to tomato, sequence motifs associated with PIF4 (E-box) or heat-responsive WRKY (W-box) transcription factors are found in *SlSPCH’s cis*-regulatory region. Under the temperature shift regime, the *SlSPCH* cis-regulatory variants again showed diverse responses. SI and SD responses in lines #1-#3 to a shift from 26°C to 34°C were slightly dampened relative to those in M82. Line *#4*, which deletes several E- and W-boxes, was essentially insensitive, and line *#5* showed a dramatic decrease in SI at 34°C (Figure 4a and S2d). The higher temperature resulted in an overall decrease in leaf area in all lines (Figure S2c).

Previous work found an inverse relationship between SD and the size of individual stomata (40) that could serve to compensate for low SD when plants were exposed to high temperature (32) We did not see this relationship in our *cis*-regulatory alleles; stomatal sizes in three mutants were similar to M82 and unaffected by temperature, and in the two mutants with significant size changes, smaller sizes were correlated with fewer stomata (Figure S2g).

We used live cell imaging and lineage tracking to determine how temperature affected stomatal lineage divisions and fates and found that the primary cellular mechanism underlying an decrease in SI was a decrease the proportion of asymmetric precursor divisions producing a stoma and pavement cell (rather than two pavement cells) at low temperature (Figure 4c). Thus, light and temperature regimes that favor low SI and SD in tomato appear to employ the same “lineage exit” strategy to do so.

Among our *SlSPCH* cis-regulatory mutants, response to changing temperature could be separated from responses to light and drought. For example, SI in line #4 was hypersensitive to light, but insensitive to temperature and drought. Line *#5* only showed a strong change in response to temperature (e.g., Figures 2a, 3a and 4a). Other *SlSPCH* cis-regulatory alleles displayed partial or dampened responses across multiple environmental conditions, indicating that different *cis*-regulatory deletions altered the magnitude and direction of stomatal responses in an allele-specific manner. Importantly, these differential responses in SI and SD were present in seedlings and in leaves of 4-week-old plants where they were linked to whole-plant traits including daily transpiration rates (Figure 2).

### Deletion of coding sequences at the SlSPCH N-terminus generates a temperature sensitive allele

The striking reduction of SI and SD at high temperature, but not in response to drought or low light, in line #5 led us to characterize in more detail the effects of the *SlSPCHmut5* allele on the sequence and function of the resultant SlSPCH protein, as well as on cellular and whole plant phenotypes (Figure 5). The *SlSPCHmut5* deletion extends into the *SlSPCH* coding region; if translation were to begin at the first in-frame Methionine codon, the resulting predicted protein would lack the basic residues of the bHLH domain but retain the helix-loop-helix domain that is highly conserved among SPCH homologues (Figure 5a). SPCH is an obligate heterodimer with partners ICE1/SCRM2, and dimerization is mediated through both the HLH domain and a C-terminal ACT domain (41). Both dimerization domains are predicted to remain intact in *SlSPCHmut5*. AlphaFold modeling of full length SlSPCH and *SlSPCHmut5*, each in complex with ICE1 on a DNA template, suggests that even missing the N-terminus, a stable dimer could form (Figure 5b-c). To assess whether plants can make a protein from the *SlSPCHmut5*variant, we created *SlSPCHmut5* reporters in a *N. benthamiana* transient expression system. We infiltrated *N. benthamiana* leaves with a plasmid co-expressing ML1p::RCI2A-mScarletI (plasma-membrane localized control for transformation) and either or *35Sp::SlSPCH-NeonGreen* (control) or *35Sp:: SlSPCHmut5-NeonGreen* at three temperatures, 22°C, 28°C and 34°C. At all temperatures, full-length *35Sp::SlSPCH-NeonGreen* was nuclear, although at higher temperature, signal was detected in fewer cells (Figure S3a, panels 1-3, quantified in S3b). At 22°C, *35Sp:: SlSPCHmut5-NeonGreen* was made and was also predominantly nuclear, though fewer transformed cells showed expression relative to the full length *SlSPCH* (Figure S3a, panel 4). At 28°C and 34°C, very little nuclear *35Sp:: SlSPCHmut5-NeonGreen* signal could be detected and most of the signal was cytosolic (Figure S3a, panels 5-6, quantified in S3b).

**Figure 5.**
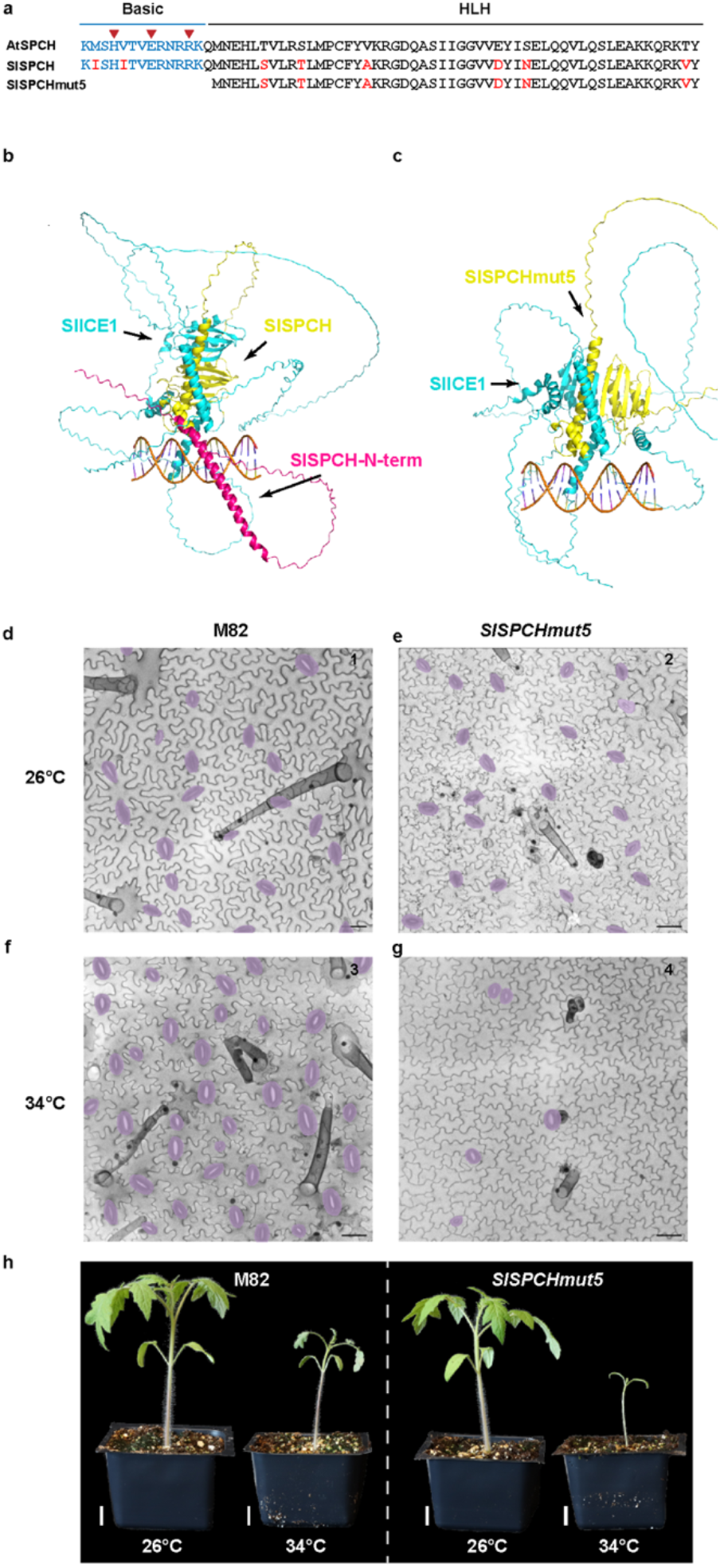
Molecular, cellular, and whole-plant characterization of a temperature-sensitive allele of *SlSPCH*. (a) Schematic of AtSPCH and SlSPCH protein sequence and predicted effect of SlSPCHmut5 allele. (b-c) AlphaFold-Multimer prediction of the SlSPCH-SlICE1 complex (b), and SlSPCHmut5-SlICE1 complex (c). SlSPCH forms a characteristic bHLH fold with partner SlICE1. A DNA-contacting alpha-helix of SlICE1 is predicted in both heterodimers, but the SlSPCHmut5 is missing a long N-terminal helix (red). (d-g) Confocal images of cotyledon epidermis in M82 (d, f) and *SlSPCHmut5* variant (e, g) at 26°C (d-e) and 34°C (f-g), stomata false-colored purple. (h) Whole plant images of M82 and *SlSPCHmut5* at specified temperatures. Scale bars in (d-g) 50 µm, (h) 20 mm.

In tomato leaves at 26°C, stomatal production and stomatal size were slightly lower in *line #5* than M82 (Fig 5d-e, S2g), but the whole plant phenotypes were indistinguishable from M82 (Figure 5h), suggesting that line #5’s stomatal performance was sufficient for normal plant function. In line #5 at 34°C, however, stomatal numbers dropped precipitously (Figure 5g) and overall growth was stunted (Figure 5h).

The loss of nuclear localization of *SlSPCHmut5* correlates with the severity of the line #5 stomatal reduction phenotype. We therefore surmise that we have generated a temperature sensitive *SlSPCH* allele whose phenotype is linked to its ability to act as a transcription factor in the nucleus. We speculate that stabilization of *SlSPCHmut5* by heterodimerization is sufficient to enable some activity at lower temperature. The *SlSPCHmut5* variant, therefore, provides a new tool to generate and test tomato lines of varying stomatal densities for engineering photosynthetic and water use efficiency phenotypes.

### Live-cell imaging reporters track the cellular responses to environmental change

The differential behaviors of edited *cis*-regulatory alleles suggested ways in *SlSPCH* could be subject to upstream regulators and to mediate environmentally responsive stomatal development. We also, however, showed tomato and Arabidopsis stomatal lineages prioritize different cell fate and division behaviors to adjust lineages in response to light and temperature (Figure 3 and 4 and (35) (42)). Thus, there remains a question of whether SPCH (and MUTE and FAMA) regulate the developmental steps in the same way in Arabidopsis and tomato. SlSPCH, SlMUTE and SlFAMA can complement loss of function mutations in their Arabidopsis mutant counterparts when expressed in Arabidopsis (43), but the expression patterns and roles of these tomato genes in their native context have not been described.

We designed native-promoter driven translational reporters and created stable transgenics in M82 (details in methods). Together, these reporters define the spatiotemporal progression of the tomato stomatal lineage (Figure 6a-c and S4a-d). Although SlSPCHpro::NeonGreen-SlSPCH (SlSPCH), SlMUTEpro::mScarlet-SlMUTE (SlSMUTE), and SlFAMApro::mTurquoise-SlFAMA (SlFAMA), follow the same general expression trends of their Arabidopsis counterparts, there are differences (Figure 6a-c and S4a-d). In newly emerging cotyledons, 0 days post emergence (dpe, Figure 6b), SlSPCH is in many epidermal cells (Figure 6c and S4a), but one day later, becomes restricted to the smaller daughters of asymmetric cell divisions (ACDs), and its expression fades in guard mother cells (GMCs; Figure 6c and S4b). SlMUTE expression starts in ACDs, peaks in GMCs and persists through the symmetric cell division (SDiv) into young guard cells (yGCs; Figure 6c and S4c). SlFAMA is expressed in young GCs (yGCs) before they form a pore (31/48 stomata) and mature GCs (mGCs) with pores (56/69 stomata) but was not detected in GMCs (0/22 GMCs) (Figure 6c and S4d). Qualitatively, SlSPCH is in the same types of asymmetrically dividing early stomatal precursor cells as AtSPCH. SlMUTE however displays broader and more prolonged expression in tomato, extending from asymmetric divisions through guard mother cells and into young GCs, in contrast to its narrow window of GMC expression in Arabidopsis (30, 31). In addition, the absence of detectable SlFAMA expression in GMCs highlights a divergence in late-lineage regulation between the two species. The lack of overt phenotypic effects on SI, SD or growth under standard growth conditions (see methods) indicates that these reporters do not measurably perturb stomatal development, establishing them as robust tools for subsequent analyses of lineage dynamics and genetic or environmental modulation of stomatal fate.

**Figure 6.**
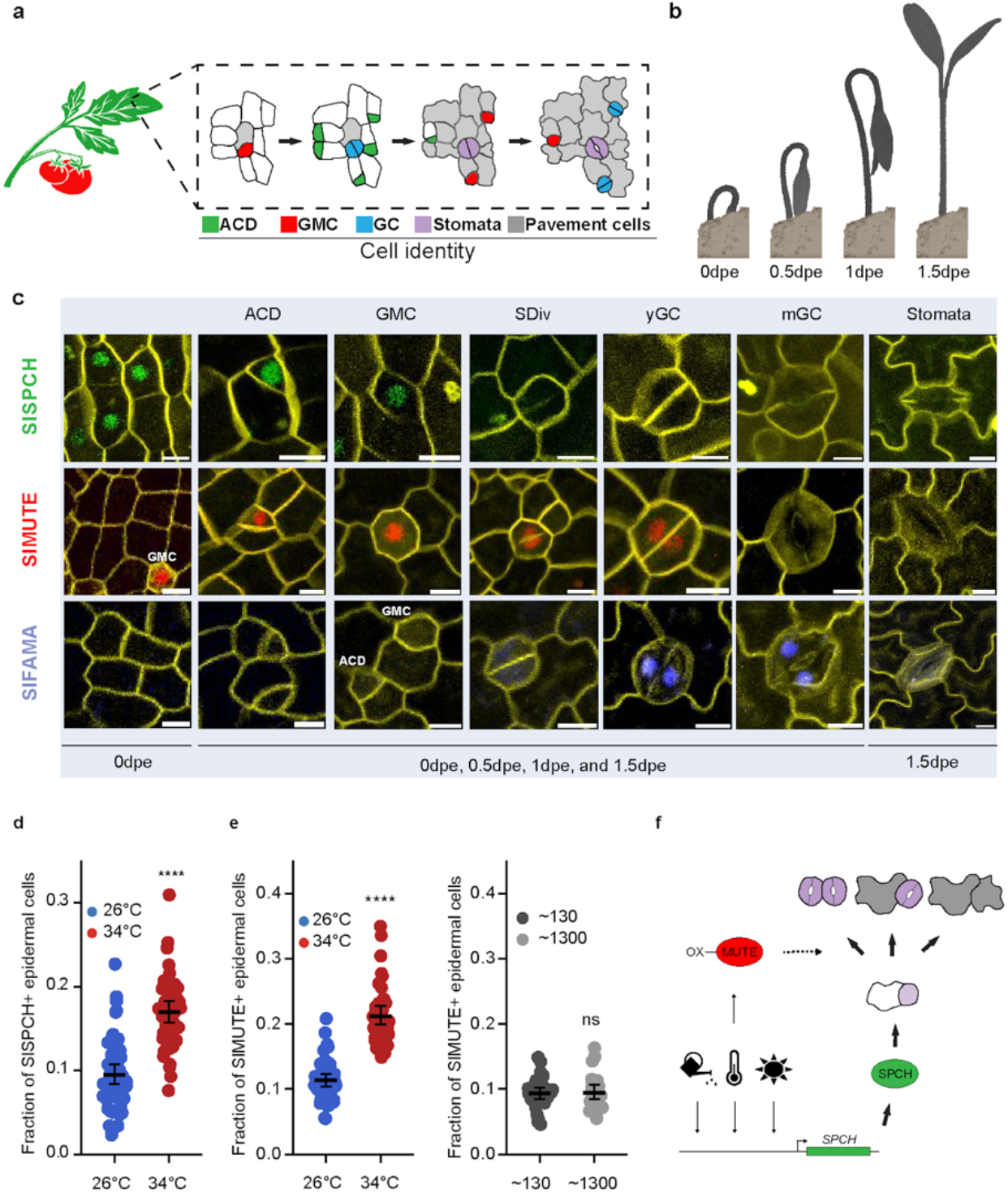
Stomatal lineage reporters provide evidence for modified lineage progression linked to capacity for environmental response. (a) Cartoon of stomatal development in tomato leaves based on observations of divisions and expression of reporters. (b) Cartoon of germination stages of tomato seedling coincident with stomatal lineage stages tracked by reporters in (c); dpe is days post-emergence. (c) Confocal images of cell type-specific expression of SlSPCH (green), SlMUTE (red) and SlFAMA (blue) reporters; colors of reporters mirror stages presented in (a). At 0dpe ACDs and a few GMCs are present. Multiple asynchronous stomatal lineages proceed toward terminal differentiation into stomata or pavement cells by 1.5dpe; individual cells are highlighted here, and larger fields of view are in Figure S4a-d. ACD-asymmetric cell division (meristemoid), GMC-guard mother cell, SDiv-symmetric division, yGC-young guard cell, and mGC-maturing guard cells. Cell outlines (yellow) visualized by propidium iodide stain. (d) Increase in fraction of SlSPCH+ epidermal cells at 34°C consistent with higher stomatal production at that temperature. (e-f) Sharp increase in fraction of SlMUTE+ epidermal cells at 34°C (e) but not in response to high light (f). (g) Summary model incorporating information from cis-regulatory *SlSPCH* alleles and stomatal lineage reporters to suggest how alteration of cell fates after SlSPCH (or SlMUTE) driven-asymmetric divisions leads to adjustment of stomatal production in response to environmental inputs.

We used SlSPCH and SlMUTE translational reporters to assess how environmental cues affect not only final SI and SD, but also the relative abundance and spatial distribution of precursor states. We focused first on temperature responses as this variable that has implications for agricultural practices in changing climates. Quantification of the SlSPCH translational reporter revealed a significant increase in the proportion of SPCH-positive cells at elevated temperature (34°C) relative to control conditions (26°C) (Figure 6d and S4e). This increase suggests that high temperature prolongs SlSPCH expression, or expands the population of cells expressing it, both of which would lead to increased stomatal production. Notably, this effect was measured as an increased fraction of reporter-positive cells rather than a change in overall epidermal cell density, indicating a specific effect on lineage allocation rather than general increase in cell proliferation.

Elevated temperature also caused a pronounced increase in the fraction of SlMUTE-reporter expressing cells (Figure 6e and S5a), accompanied by a substantial rise in stomatal production (Figure S5b). High temperature also resulted in frequent violations of the one-cell spacing rule, producing clusters of adjacent stomata (Figure S5c-d), whereas clustering was rare at 26°C (Figure S5c-d). Stomatal clustering was not observed in the SlSPCH reporter line at any temperature (Figure S4), nor did SlMUTE induce stomatal clusters in response to higher light (Figure S5b-c, e), despite our light shift regime inducing an increase in SI at least as great as our temperature shift regime in M82 controls (Figure 3a-b, 4a and S5b). Thus, increased stomatal production *per se* is not sufficient to induce clustering in SlMUTE reporter lines, suggesting temperature-specific interaction between SlMUTE expression and activity and lineage control mechanisms.

## DISCUSSION

Plants perceive and interpret environmental cues and effect developmental changes. Understanding the signaling, transcriptional and cellular responses that bridge input and output can reveal fundamental mechanisms of information flow and provide materials for creating climate-resilient crops. Here, using CRISPR/Cas9-enabled genome editing of the *cis-*regulatory region of the stomatal lineage regulatory factor *SPCH* in tomato, we showed that stomatal production could be made more or less sensitive to light, drought and temperature cues. Importantly, these sensitivities were allele-specific, with some *cis*-regulatory deletions selectively affecting responsiveness to a single environmental input while leaving others intact. *SlSPCHmut5*, an allele whose strong temperature-sensitive activity response appears linked to the subcellular localization of the protein provides both insights into intrinsic regulation of SPCH and a tool for manipulating stomatal production. By combining molecular data on alleles with reporters and cellular tracking data, we identified a cell fate switching mechanism that underlies adjustments made in response to light and to temperature. These SlSPCH alleles and lines bearing reporters for SlSPCH, SlMUTE and SlFAMA comprise a genetic toolkit that will enable finer dissection of stomatal lineage behaviors under many conditions and in other mutant backgrounds.

Cis-regulatory elements dictate cell-type specific expression and/or can tune expression in response to the environment. We showed that deletions of the 5’ cis-regulatory region of *SlSPCH* result in altered stomatal response to environmental change while leaving overall stomatal patterning (e.g. spacing) intact, suggesting that we identified mainly tuning elements. This outcome was by design, given our initial screen was for CRISPR/Cas9-edited plants with overall wildtype appearance under standard growth conditions. Consistent with this, most alleles displayed partial or dampened responses rather than complete loss of environmental responsiveness, indicating quantitative modulation rather than binary control. Within the 3Kb SlSPCH 5’region we targeted, there are predicted sites for the light response-mediating transcription factor HY5 (including A-boxes), two bHLH target sites (E-Boxes) of the type used by the temperature-mediating transcription factor PIF4, and abscisic acid response elements (ABREs) as well as drought responsive MSBPs and DRE/CRTs. Line #4, which was insensitive to temperature, but hypersensitive to light, disrupts HY5 and two E-boxes (Figure 1b and S1b). Line #3, a relatively small deletion which exhibits a dampened light response, interrupts one of the A-boxes (Figure 1b and S1b). Future experiments may reveal the roles of tomato HY5 and PIF genes (44, 45) in stomatal regulation. It will be particularly interesting if HY5 is a direct transcriptional regulator of *SlSPCH* because in Arabidopsis, there is an intermediary between HY5 and regulation of SPCH protein (4). Sequence predictions in the *SlSPCH* upstream region, and alignment with SPCH 5’ regions from other Solanaceae and from Arabidopsis reveal a number of conserved regions (CNS, Figure S1c). Some of the Solanaceae CNS are included in our characterized deletion lines (Figure 1b and S1b) but the CNS conserved between tomato and Arabidopsis (red in Figure S1c) lies further upstream. Generation of additional cis-regulatory alleles that specifically target CNS may be a productive strategy to identify additional regulatory elements, including both those that drive cell-type specific expression and those that affect *SlSPCH* production in response to additional environmental inputs.

The behavior of the *SlSPCHmut5* allele that eliminates the N-terminal region, including the predicted DNA-binding region of the bHLH domain, challenges our assumptions about which regions of the SlSPCH protein are essential, and for which activities. At 26°C line #5 produced fewer and slightly smaller stomata than M82 (Figure S2g), but the stomata appear morphologically normal and well-patterned (Figure 5d-e), therefore the *SlSPCHmut5* variant must be capable of performing the general SlSPCH functions of inducing asymmetric cell divisions and promoting eventual GC identity. At elevated temperature, *SlSPCHmut5* confers a consistent and specific defect in stomatal production and size, consistent with generation of a less functional protein. Interestingly, our untargeted mutagenesis mimics the results reported in work designed to generate hypomorphic or conditional alleles by removing the N-terminus and forcing use of an internal Methionine (46). Although it may seem surprising to ascribe transcriptional regulator activity to a bHLH missing the motif that typically mediates DNA contacts, there are classes of transcription factors that can function without sequences N-terminal of the HLH domains (47), and previous deletion and domain swap experiments showed that AtMUTE, and to a lesser extent, AtSPCH, can function when three putative DNA-binding residues are replaced (48). It was previously suggested that in heterodimer pairs of SPCH or MUTE with ICE1, it is ICE1 that makes contact with DNA, and SPCH provides some other essential activities for transcriptional activation (49). Loss of the SlSPCH N-terminal region is also predicted to eliminate a nuclear localization signal, and we see that, at high temperatures, *SlSPCHmut5* cannot be retained in the nucleus (Figure S3a). That *SlSPCHmut5* is ever nuclear we attribute to its heterodimerization with *SlICE* (Figure 5c). We speculate that, given ICE1’s activity promoting gene expression at low temperatures (50, 51), it may be less present or active at warm ones, and low ICE1 would be insufficient to retain the N-terminal deleted SlSPCH.

Ultimately how do the manipulations we made to the *SlSPCH* locus lead to environment-dependent changes in stomatal production? Cell tracking data in M82 suggest that SlSPCH-induced ACDs normally resolve into either a stoma and pavement cell or two pavement cells, and light and temperature modulate SlSPCH production and/or activity to tip this balance. One hypothesis is that quantitative shift in *SlSPCH* in cells about to undergo ACDs defines whether one or no cells acquire stomatal fate after ACD. We, however, prefer a second explanation that also considers the broadened expression pattern of SlMUTE and the effects we saw in SlMUTE translational reporter lines (Figure 6g). Overexpression of MUTE throughout Arabidopsis leads to the formation of extra stomata by inducing non-stomatal cells to take on stomatal fate (52). The ectopic stomata produced by the extra copy of *SlSMUTE*, however, are morphologically normal, and the orientation of the GC pairs suggests that they do not originate from extra divisions of GCs (Figure S5d, compared to (53)). Therefore, we hypothesize that the extra SlMUTE dosage is revealing more of the range in the post-ACD toggle between pavement cell and GC identity. Combined with the larger overlap in SlSPCH and SlMUTE expression in tomato, this suggests that tomato ACDs produce a continuum of cell fates. Under conditions optimal for productive photosynthesis and leaf growth, ACDs result in one GC and one pavement cell. The ACDs are induced by SlSPCH, but the resulting choice of fate is dictated by SlSPCH targets (or SlSPCH in complex with a target) and among those targets is *SlMUTE*. Under limiting environmental conditions, SlSPCH levels or activity are insufficient to induce fate-promoting targets, and both products of an ACD become pavement cells. In high temperature conditions, elevated SlSPCH and/or its direct targets would have both endogenous and transgene-derived SlMUTE to induce and this higher expression would push the ACDs to yield two stomata. Alternatively, SlMUTE itself could be temperature responsive and an additional copy of SlMUTE could cause SlMUTE to accumulate to levels high enough to pass a cell fate threshold in both daughters of an ACD.

Nearly all future climate predictions suggest that temperatures and atmospheric [CO_2_] will increase, and severe weather events (drought, flooding) will become more prevalent (54, 55). Stomata, with their roles in capturing atmospheric carbon and regulating plant transpiration, which has both water transport and leaf cooling components, must integrate and prioritize potentially conflicting signals to maintain plant health. While the increase in stomatal production in response to increasing light intensity appears to occur in most plants, the response to increased temperature can vary, as seen comparing tomato and Arabidopsis, and even among Arabidopsis accessions. Our data further show that this variation can be genetically uncoupled at the level of a single developmental regulator. These diverging responses to warm temperature may reflect different priorities for conserving water and leaf cooling. It is therefore hard to predict, but important to test, the combinatorial effects of predicted future climate conditions on plants. Genetic tools that can alter sensitivity to specific inputs, such as the SlSPCH lines generated in this work, will be instrumental in deciphering complex response and may also be useful to assay the effects of having tomato plants with different stomatal numbers for growth in large-scale or urban agricultures systems of the future.

## Author Contributions

IN designed research, performed research, analyzed data, and wrote the paper; AB, DS, HP, EPZ, YK, and NKS performed research, JE performed research and analyzed data, YT analyzed data, DCB designed research, analyzed data and wrote the paper.

## REFERENCES

1. C. B. Engineer, et al., CO2 Sensing and CO2 Regulation of Stomatal Conductance: Advances and Open Questions. Trends Plant Sci. 21, 16–30 (2016).

2. J. C. McElwain, M. Steinthorsdottir, Paleoecology, Ploidy, Paleoatmospheric Composition, and Developmental Biology: A Review of the Multiple Uses of Fossil Stomata. Plant Physiol. 174, 650–664 (2017).

3. P. L. M. Lang, et al., Century-long timelines of herbarium genomes predict plant stomatal response to climate change. Nat. Ecol. Evol. 8, 1641–1653 (2024).

4. S. Wang, et al., Light regulates stomatal development by modulating paracrine signaling from inner tissues. Nat. Commun. 12, 3403 (2021).

5. M. F. Pompelli, S. C. V. Martins, E. F. Celin, M. C. Ventrella, F. M. Damatta, What is the influence of ordinary epidermal cells and stomata on the leaf plasticity of coffee plants grown under full-sun and shady conditions? Braz. J. Biol. 70, 1083–1088 (2010).

6. O. S. Lau, et al., Direct Control of SPEECHLESS by PIF4 in the High-Temperature Response of Stomatal Development. Curr. Biol. 28, 1273-1280.e3 (2018).

7. Y. Zheng, et al., Effects of experimental warming on stomatal traits in leaves of maize (Zea may L.). Ecol. Evol. 3, 3095–3111 (2013).

8. H. I. U. Caldera, W. A. J. M. De Costa, F. I. Woodward, J. A. Lake, S. M. W. Ranwala, Effects of elevated carbon dioxide on stomatal characteristics and carbon isotope ratio of Arabidopsis thaliana ecotypes originating from an altitudinal gradient. Physiol. Plant. 159, 74–92 (2017).

9. H. Wang, et al., BZU2/ZmMUTE controls symmetrical division of guard mother cell and specifies neighbor cell fate in maize. PLoS Genet. 15, e1008377 (2019).

10. M. T. Raissig, et al., Mobile MUTE specifies subsidiary cells to build physiologically improved grass stomata. Science 355, 1215–1218 (2017).

11. M. T. Raissig, E. Abrash, A. Bettadapur, J. P. Vogel, D. C. Bergmann, Grasses use an alternatively wired bHLH transcription factor network to establish stomatal identity. Proc. Natl. Acad. Sci. U. S. A. 113, 8326–8331 (2016).

12. Z. Jiao, et al., PdEPFL6 reduces stomatal density to improve drought tolerance in poplar. Ind. Crops Prod. 182, 114873 (2022).

13. M. Clemens, et al., VvEPFL9-1 Knock-Out via CRISPR/Cas9 Reduces Stomatal Density in Grapevine. Front. Plant Sci. 13, 878001 (2022).

14. C. C. C. Chater, R. S. Caine, A. J. Fleming, J. E. Gray, Origins and Evolution of Stomatal Development. Plant Physiol. 174, 624–638 (2017).

15. R. S. Caine, et al., Rice with reduced stomatal density conserves water and has improved drought tolerance under future climate conditions. New Phytol. 221, 371–384 (2019).

16. C. C. Chater, et al., Origin and function of stomata in the moss Physcomitrella patens. Nat Plants 2, 16179 (2016).

17. K. H. McKown, D. C. Bergmann, Stomatal development in the grasses: lessons from models and crops (and crop models). New Phytol. 227, 1636–1648 (2020).

18. L. R. Lee, D. C. Bergmann, The plant stomatal lineage at a glance. J. Cell Sci. 132 (2019).

19. L. C. Chua, O. S. Lau, Stomatal development in the changing climate. Development 151 (2024).

20. D. Samakovli, et al., YODA-HSP90 Module Regulates Phosphorylation-Dependent Inactivation of SPEECHLESS to Control Stomatal Development under Acute Heat Stress in Arabidopsis. Mol. Plant 13, 612–633 (2020).

21. C. Han, et al., TOR and SnRK1 fine tune SPEECHLESS transcription and protein stability to optimize stomatal development in response to exogenously supplied sugar. New Phytol. 234, 107–121 (2022).

22. C. Klermund, et al., LLM-Domain B-GATA Transcription Factors Promote Stomatal Development Downstream of Light Signaling Pathways in Arabidopsis thaliana Hypocotyls. Plant Cell 28, 646–660 (2016).

23. X. Yang, L. G. S, Z. Zhou, D. Urano, O. S. Lau, Abscisic acid regulates stomatal production by imprinting a SnRK2 kinase–mediated phosphocode on the master regulator SPEECHLESS. Science Advances 8, eadd2063 (2022).

24. D. Rodríguez-Leal, Z. H. Lemmon, J. Man, M. E. Bartlett, Z. B. Lippman, Engineering Quantitative Trait Variation for Crop Improvement by Genome Editing. Cell 171, 470-480.e8 (2017).

25. D. Rodriguez-Leal, et al., Evolution of buffering in a genetic circuit controlling plant stem cell proliferation. Nat. Genet. 51, 786–792 (2019).

26. A. Hendelman, et al., Conserved pleiotropy of an ancient plant homeobox gene uncovered by cis-regulatory dissection. Cell 184, 1724-1739.e16 (2021).

27. C. A. MacAlister, K. Ohashi-Ito, D. C. Bergmann, Transcription factor control of asymmetric cell divisions that establish the stomatal lineage. Nature 445, 537–540 (2007).

28. S. A. M. McAdam, T. J. Brodribb, Linking turgor with ABA biosynthesis: Implications for stomatal responses to vapor pressure deficit across land plants. Plant Physiol. 171, 2008–2016 (2016).

29. Y. Tanaka, T. Nose, Y. Jikumaru, Y. Kamiya, ABA inhibits entry into stomatal-lineage development in Arabidopsis leaves. Plant J. 74, 448–457 (2013).

30. Y. Uno, et al., Arabidopsis basic leucine zipper transcription factors involved in an abscisic acid-dependent signal transduction pathway under drought and high-salinity conditions. Proc. Natl. Acad. Sci. U. S. A. 97, 11632–11637 (2000).

31. T. Urao, K. Yamaguchi-Shinozaki, S. Urao, K. Shinozaki, An Arabidopsis myb homolog is induced by dehydration stress and its gene product binds to the conserved MYB recognition sequence. Plant Cell 5, 1529–1539 (1993).

32. M. L. Pérez-Bueno, et al., An extremely low stomatal density mutant overcomes cooling limitations at supra-optimal temperature by adjusting stomatal size and leaf thickness. Front. Plant Sci. 13, 919299 (2022).

33. N. Zoulias, J. Brown, J. Rowe, S. A. Casson, HY5 is not integral to light mediated stomatal development in Arabidopsis. PLoS One 15, e0222480 (2020).

34. A. Vatén, C. L. Soyars, P. T. Tarr, Z. L. Nimchuk, D. C. Bergmann, Modulation of Asymmetric Division Diversity through Cytokinin and SPEECHLESS Regulatory Interactions in the Arabidopsis Stomatal Lineage. Dev. Cell 47, 53-66.e5 (2018).

35. I. Nir, et al., Evolution of polarity protein BASL and the capacity for stomatal lineage asymmetric divisions. Curr. Biol. 32, 329-337.e5 (2022).

36. J. Zhu, et al., Effect of simulated warming on leaf functional traits of urban greening plants. BMC Plant Biol. 20, 139 (2020).

37. Y. Li, et al., SlSnRK2.3 interacts with SlSUI1 to modulate high temperature tolerance via Abscisic acid (ABA) controlling stomatal movement in tomato. Plant Sci. 321, 111305 (2022).

38. J. Urban, M. W. Ingwers, M. A. McGuire, R. O. Teskey, Increase in leaf temperature opens stomata and decouples net photosynthesis from stomatal conductance in Pinus taeda and Populus deltoides x nigra. J. Exp. Bot. 68, 1757–1767 (2017).

39. K. Gasparini, et al., The Lanata trichome mutation increases stomatal conductance and reduces leaf temperature in tomato. J. Plant Physiol. 260, 153413 (2021).

40. H. Dittberner, et al., Natural variation in stomata size contributes to the local adaptation of water-use efficiency in Arabidopsis thaliana. Mol. Ecol. 27, 4052–4065 (2018).

41. H. Seo, et al., Intragenic suppressors unravel the role of the SCREAM ACT-like domain for bHLH partner selectivity in stomatal development. Proc. Natl. Acad. Sci. U. S. A. 119 (2022).

42. M. Rath, N. K. Sharma, M. Mani, D. C. Bergmann, Stomatal setpoints and environmental responsiveness are sculpted by developmental trajectories. bioRxiv 2025.12.22.696041 (2025).

43. A. Ortega, A. de Marcos, J. Illescas-Miranda, M. Mena, C. Fenoll, The Tomato Genome Encodes SPCH, MUTE, and FAMA Candidates That Can Replace the Endogenous Functions of Their Arabidopsis Orthologs. Front. Plant Sci. 10, 1300 (2019).

44. D. Rosado, et al., Phytochrome Interacting Factors (PIFs) in Solanum lycopersicum: Diversity, Evolutionary History and Expression Profiling during Different Developmental Processes. PLoS One 11, e0165929 (2016).

45. C. Zhang, et al., Pivotal roles of ELONGATED HYPOCOTYL5 in regulation of plant development and fruit metabolism in tomato. Plant Physiol. 189, 527–540 (2022).

46. M. Yoshimura, T. Mamiya, N. Takahashi, T. Ishida, A start codon-targeted genome editing strategy for generating hypomorphic mutants of lethal plant genes. Plant Biotechnol. (Tsukuba) 42, 509–512 (2025).

47. Y. Hao, X. Zong, P. Ren, Y. Qian, A. Fu, Basic Helix-Loop-Helix (bHLH) Transcription Factors Regulate a Wide Range of Functions in Arabidopsis. Int. J. Mol. Sci. 22 (2021).

48. K. A. Davies, D. C. Bergmann, Functional specialization of stomatal bHLHs through modification of DNA-binding and phosphoregulation potential. Proc. Natl. Acad. Sci. U. S. A. 111, 15585–15590 (2014).

49. A. Liu, et al., Cell Fate Programming by Transcription Factors and Epigenetic Machinery in Stomatal Development. bioRxiv 2023.08.23.554515 (2023).

50. T. Ma, et al., Arabidopsis LFR, a SWI/SNF complex component, interacts with ICE1 and activates ICE1 and CBF3 expression in cold acclimation. Front. Plant Sci. 14, 1097158 (2023).

51. R. Lin, et al., CALMODULIN6 negatively regulates cold tolerance by attenuating ICE1-dependent stress responses in tomato. Plant Physiol. 193, 2105–2121 (2023).

52. L. J. Pillitteri, D. B. Sloan, N. L. Bogenschutz, K. U. Torii, Termination of asymmetric cell division and differentiation of stomata. Nature 445, 501–505 (2007).

53. L. B. Lai, et al., The Arabidopsis R2R3 MYB proteins FOUR LIPS and MYB88 restrict divisions late in the stomatal cell lineage. Plant Cell 17, 2754–2767 (2005).

54. S. Alamos, P. M. Shih, How to engineer the unknown: Advancing a quantitative and predictive understanding of plant and soil biology to address climate change. PLoS Biol. 21, e3002190 (2023).

55. M. Haworth, G. Marino, F. Loreto, M. Centritto, Integrating stomatal physiology and morphology: evolution of stomatal control and development of future crops. Oecologia 197, 867–883 (2021).

