## Supplemental Methods, Figures S1-5; Tables S2-4 for "Genetic encoding of climate-responsive stomatal developmental plasticity in tomato"

**This PDF file includes:**

Supporting Materials and Methods
Supporting References
Supplemental Figures S1- S5
Supplemental Tables S2-S4

**Other supporting materials for this manuscript include the following:**

Table S1

### METHODS

Plant growth: Tomato plants (*Solanum lycopersicum* cv. M82) were grown in a Percival chamber model CU22L, or Percival growth room, model AR-1015L3 set to a photoperiod of 16 hr light/8 hr dark, light intensity of approximately 250 µmol m2 s-1, and 26 °C or 34 °C. For propagation, plants were grown in a greenhouse under natural day length conditions, at 700–1200 µmol m2 s-1 and 18 – 29 °C. For light experiments the Percival AR-1015L3 growth room light intensity set to 130 µmol $m^2$ $s^{-1}$ for low light experiment and 1300 µmol $m^2$ $s^{-1}$ for high light experiment.

Tomato DNA constructs and plant transformation: For tomato SlSPCH, SlMUTE, and SlFAMA marker lines, the promoters and CDS (primers used for cloning are in Table S3) were cloned into a level 0 MoClo part using the Golden Gate cloning system (1) and then fused to NeonGreen, mScarletI and mTurquoise respectively with a NOS terminator to form a level 1 construct. Each level 1 was transferred to a level 2, together with a kanamycin resistance cassette. The constructs were sub-cloned into the pAGM4723 binary vector and were introduced into *Agrobacterium tumefaciens* strain GV3101 by electroporation. The constructs were transferred to M82 cotyledons, using transformation and regeneration methods described in (2). Kanamycin-resistant T0 plants were grown and at least four independent transgenic lines were selected and self-pollinated to generate homozygous transgenic lines for each reporter line.

Tomato SPCH promoter CRISPR/Cas9 mutagenesis, plant transformation, and selection of mutant alleles 11 single-guide RNAs (sgRNAs) spanning 3000 bases upstream of the TSS of SlSPCH were designed using the CRISPR-P tool (3). Primers used for sgRNAs and subsequent genotyping are in Table S4. The gRNAs and promoter were assembled using the Golden Gate cloning system as described in (1). The final binary vector including zCas9, the gRNAs and NPTII, assembled in pAGM4723, was introduced into *Agrobacterium tumefaciens* strain GV3101 by electroporation. The construct was transferred into M82 cotyledons using transformation and regeneration methods described by (2). T0 transgenic plants resistant to Kanamycin were grown and independent lines were selected and self-pollinated to generate homozygous lines. For genotyping of the transgenic lines, genomic DNA was extracted, and each plant was genotyped by PCR for the presence of the zCas9. The positive lines for the zCas9 were further genotyped for mutations in SlSPCH promoter (Solyc03g007410) using a forward primer 3.1Kb upstream to the ATG and a reverse primer 155 bp downstream to the ATG, these pair primers cover the 11 gRNAs

Microscopy, image analysis and processing The fluorescence imaging experiments on the tomato transgenic plants were performed on either a Leica SP5, SP8 or Stellaris confocal microscope with HyD detectors using 25X NA0.95 and 40x NA1.1 water objective with image size 1024*1024 and digital zoom from 1x to 2x. To quantify the stomatal index of the tomato *SlSPCH* cis-regulatory mutants, 14 dpe cotyledons were imaged and the fraction of stomata/total epidermal cells was computed from regions of approximately 0.4 mm2. To calculate SD, stomata were counted across the entire field of view of approximately 0.4 mm2. Stomatal length is the long-axis of the stomatal complex measured at the interface of two guard cells in mature complexes with pores. Still or time-course images of ML1p::RCI2A-NeonGreen, propidium iodide (PI), and FM4-64 fluorescence in tomato were obtained from a Leica Stellaris confocal microscope with HyD detectors using 25x water objective with image size 1024*1024 and digital zoom from 1x to 2x. Time-course experiments on tomato cotyledons were done as described by (4). All raw fluorescence image Z-stacks were projected with STD Slices in FIJI.

Drought experiment M82 controls and the *SlSPCH* cis-regulatory mutants were grown under controlled conditions of sufficient water (100%) or water deficiency of 30% relative soil water content (RSWC) for three weeks. All plants were then grown under a sufficient water regime for a one-week recovery period. We used a high-throughput multi-lysimeter system (8) to monitor and control the water content of the pots, and to measure whole plant transpiration, and canopy conductance. With each pot placed on a load cell with digital output, we were able to precisely control water levels and analyze plant water loss and related physiological traits. We measured the daily whole-plant transpiration as described in (8) and the SI of expanded 5th leaves in the four-week-old plants. SI was measured on regions of approximately 0.4 $mm^2$

Transient expression in *Nicotiana benthamiana*: *N. benthamiana* plants were grown for six weeks in a phytotron greenhouse under natural day length conditions, at 700–1200 mmol m-2 s-1, with daytime temperature of 22°C, then moved to 22°C, 28°C, and 34°C for Agrobacterium infiltration and subsequent analysis of protein expression as indicated. Plants were at least 6 weeks old at time of experiment. For transient expression, dual expression constructs with expression control ML1pro::RCI2A-mScarlet-I and either 35Spro:SlSPCH-NeonGreen or 35Spro:SLSPCHmut5-NeonGreen were cloned using the Golden Gate system (1) into the binary vector pAGM4723 with rbcS terminator for SPCH variants, and NOS terminator for control. The binary vectors were introduced into *Agrobacterium tumefaciens* strain GV3101 and infiltrated into mature tobacco leaves as described in (5).

The analysis of stomatal density and index responses to changes in temperature in Arabidopsis of different accessions, followed growth condition and temperature shift protocols in (6). Seeds for accessions Col-0 (CS22625), Bur-0 (CS77833) and Kz-9 (CS22607) were obtained from the ABRC, Ohio State University, USA. Tissues were cleared by placing seedlings in 7:1 Ethanol:Acetic acid for 3 days, rinsing in 1M Potassium hydroxide for 30 minutes, then placing in water before mounted with Hoyer's solution. DIC images of the abaxial surface were collected at 20X using a Leica-DFC9000GTC-VSC11962 with a 1.623 $mm^2$ field of view and 50 µm z-step size. To calculate SD, stomata were counted across the entire field of view. To calculate SI, stomata and other epidermal cells were counted on a 0.370 $mm^2$ leaf blade region selected to avoid lying over major veins. Cells were counted using ImageJ with the CellCounter plugin. Results were plotted using RStudio with the "tidyverse" suite of packages.

Gene identifiers: *S. lycopersicum* v.M82: SlSPCH (Solyc03g007410); SlMUTE (Solyc1g080050); SlFAMA (Solyc05g05366010) *A. thaliana:* SPCH (At5g53210), MUTE (At3g06120) FAMA (At3g24140)

Quantification and statistical analyses

All statistical analyses in this manuscript were performed in RStudio (R Development Core Team., 2020). Statistical parameters for each analysis are indicated in the figure legends. For significance testing, unpaired Mann-Whitney *U* tests were conducted with the wilcox_test function from the rstatix package (7). Bonferroni corrections were performed when more than 2 pair-wise comparisons were conducted, and Bonferroni corrected p-values are indicated in all figures. Additional statistical details are in Table S2.


1. E. Weber, R. Gruetzner, S. Werner, C. Engler, S. Marillonnet, Plos One (2011).

2. S. McCormick, "Transformation of tomato with Agrobacterium tumefaciens" in *Plant Tissue Culture Manual*, (Springer, 1991), pp. 311–319.

3. Y. Lei, *et al.*, CRISPR-P: a web tool for synthetic single-guide RNA design of CRISPR-system in plants. *Mol. Plant* **7**, 1494–1496 (2014).

4. I. Nir, *et al.*, Evolution of polarity protein BASL and the capacity for stomatal lineage asymmetric divisions. *Curr. Biol.* **32**, 329–337.e5 (2022).

5. Y. Zhang, P. Wang, W. Shao, J.-K. Zhu, J. Dong, The BASL polarity protein controls a MAPK signaling feedback loop in asymmetric cell division. *Dev. Cell* **33**, 136–149 (2015).

6. O. S. Lau, *et al.*, Direct Control of SPEECHLESS by PIF4 in the High-Temperature Response of Stomatal Development. *Curr. Biol.* **28**, 1273–1280.e3 (2018).

7. A. Kassambara, rstatix: Pipe-friendly framework for basic statistical tests. R package version 0.7. 0 (2021).

8. O. Halperin, A. Gebremedhin, R. Wallach, M. Moshelion, High-throughput physiological phenotyping and screening system for the characterization of plant-environment interactions. *Plant J.* **89**, 839–850 (2017).

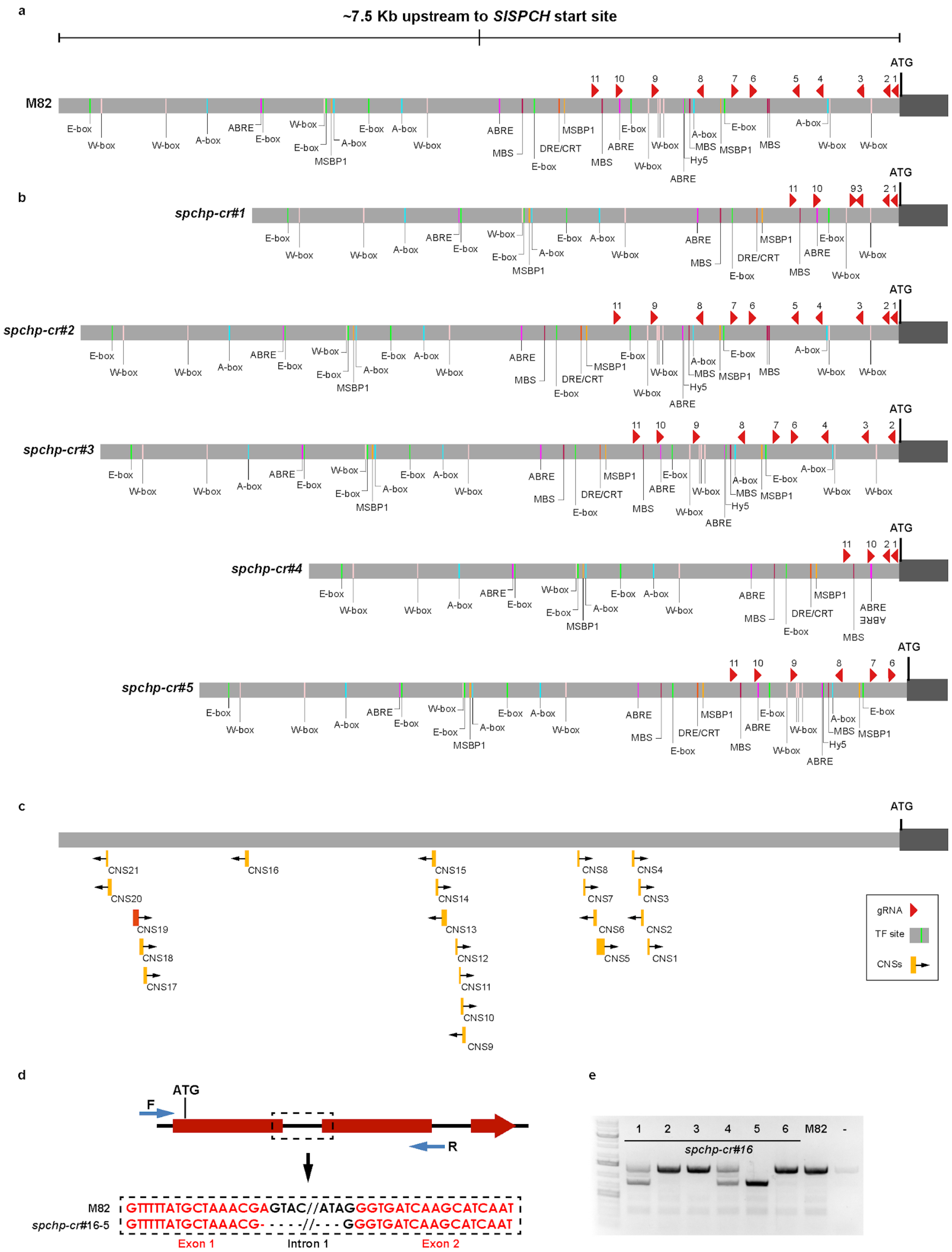
a
~7.5 Kb upstream to SlSPCH start site
ATG
M82
11 10 9 8 7 6 5 4 3 2 1
E-box W-box A-box ABRE MSBP1 MBS DRE/CRT Hy5
b
spchp-cr#1
spchp-cr#2
spchp-cr#3
spchp-cr#4
spchp-cr#5
c
CNS21 CNS20 CNS19 CNS18 CNS17 CNS16 CNS15 CNS14 CNS13 CNS12 CNS11 CNS10 CNS9 CNS8 CNS7 CNS6 CNS5 CNS4 CNS3 CNS2 CNS1
gRNA
TF site
CNSs
d
ATG
F
R
M82 GTTTTATGCTAAACGAGTAC//ATAGGGTGATCAAGCATCAAT
spchp-cr#16-5 GTTTTATGCTAAACG- - - - -//- - - GGGTGATCAAGCATCAAT
Exon 1
Intron 1
Exon 2
e
1 2 3 4 5 6 M82 -
spchp-cr#16

**Figure S1 Potential regulatory elements in M82 *SlSPCH* promoter compared with CRISPR-Cas9 generated deletion alleles**

(a-b) Annotation of the *SlSPCH cis*-regulatory region with gRNAs (red triangles) and predicted transcription factor (TF) binding sites in M82 (a) and in five CRISPR-Cas9 generated alleles characterized in detail in this paper (b). (c) Conserved Non-Coding Sequences (CNS) found between *S. lycopersicum* and other plant species. Orange CNSs are conserved only within the Solanaceae family, and the red CNS (CNS19) is conserved among eudicots including Arabidopsis. (d) Description of mutation in *SlSPCH* line #16 (phenotype in Figure 1). This putative *SlSPCH* null allele (frameshift in coding region) has a deletion (dashed square) more than 300 bp away from gRNAs. The black arrow points to the exon (red) and intron (black) sequences that are deleted (e) PCR genotyping of offspring from plant heterozygous for the #16 allele showing segregation: plants 1 and 4 are heterozygotes, 5 is a homozygous mutant and plants 2, 3 and 6 are wild type at the *SlSPCH* locus.

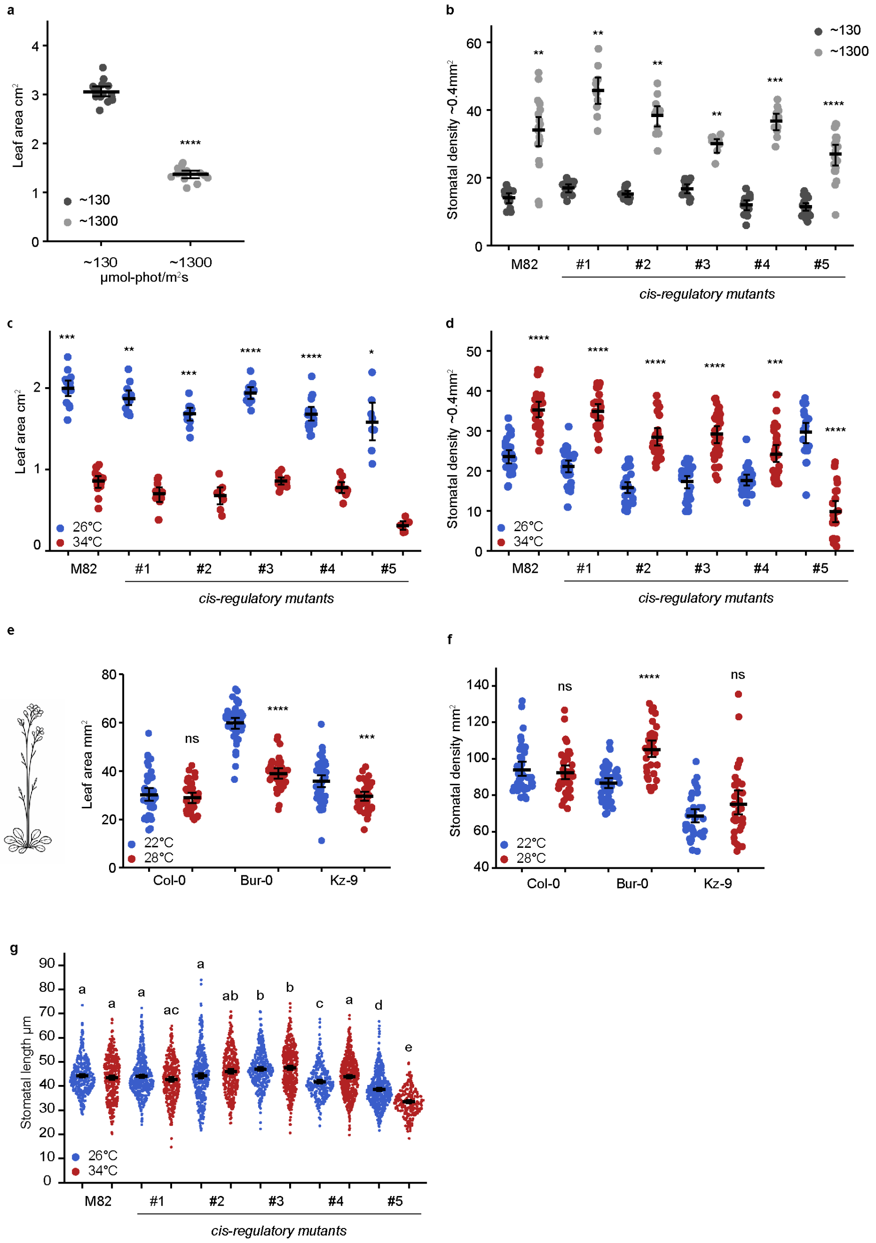


**Figure S2 Additional SI, SD, stomatal morphology, and leaf area measurements of tomato and Arabidopsis variants in response to changing environmental conditions (related to Figures 2, 3 and 4)**

(a) Plot of leaf area changes in M82 in response to light. n = 13 leaves. (b) Plot of stomatal density changes in M82 and *SlSPCH* mutants in response to changes in light intensity. Note the data in this plot are identical to those in Figure 2b, they have been repeated here in slightly modified form for ease of comparison. (c) Plot of in M82 and *SlSPCH* mutant leaf area changes in response to changes in temperature. (d) Plot of stomatal density changes in M82 and *SlSPCH* mutants in response to changes in temperature. (e) Plot of leaf area changes in Arabidopsis accessions in response to changes in temperature. (f) Plot of stomatal density changes in Arabidopsis accessions in response to changes in temperature. (g) Plot of stomatal length in M82 and *SlSPCH* mutants in response to temperature. Statistical tests in all plots are represented as mean ± 95% confidence interval. Bonferroni-corrected p values from Mann-Whitney U test are *P < 0.05; **P < 0.01; ***P < 0.001; ****P < 0.0001. n.s.: P > 0.05, not significant. Sample sizes in (a) n = 13 leaves., in (b) n = 8-21 fields from 3-5 cotyledons, in (C) n = 8-18 leaves, in (d) n = 20-30 0.4 mm$^2$ fields from 4-6 cotyledons, for (e) and (f), each point represents an individual plant (n= >34). Sample sizes in (g) n = 20-30 0.4 mm2 fields from 4-6 cotyledons, each point represents an individual stoma (n = 100-500).

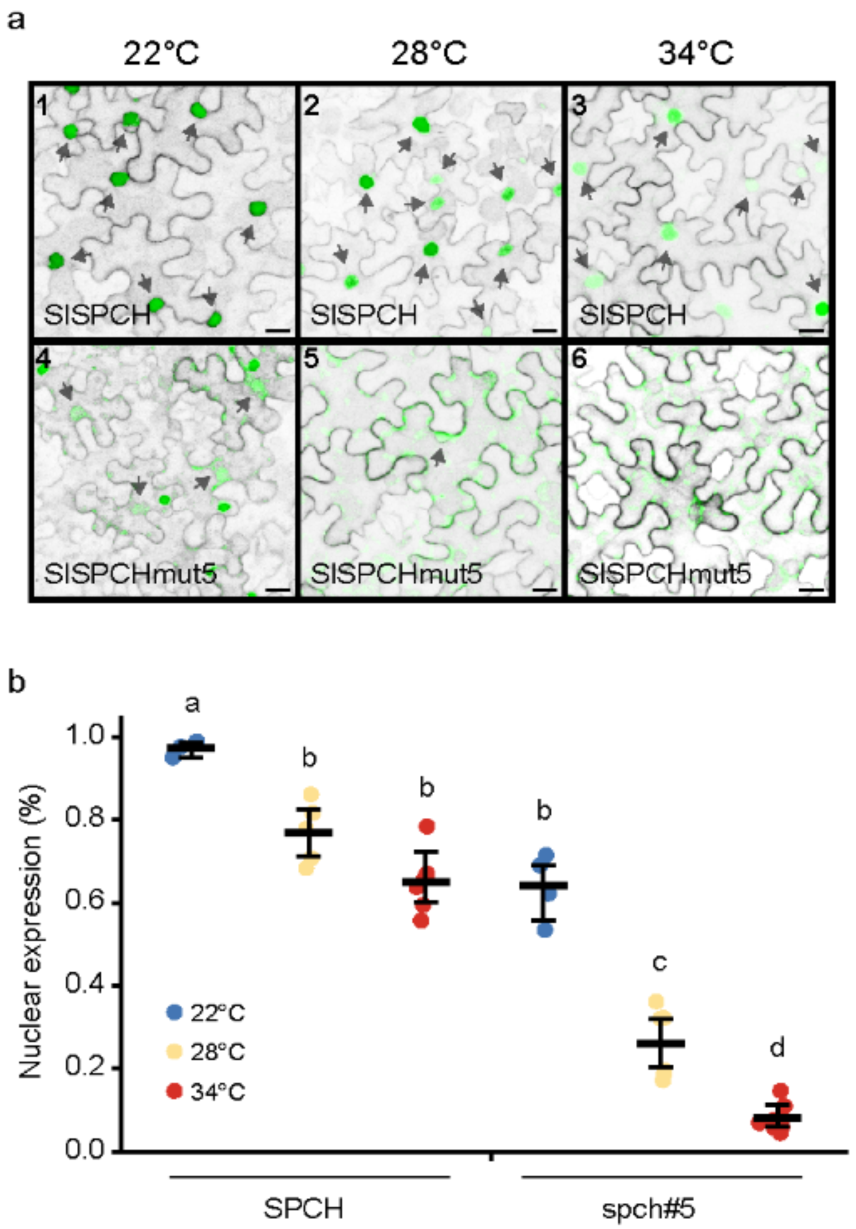


**Figure S3 Enhanced loss of nuclear localization in response to high temperature in SlSPCH reporter recapitulating *SlSPCHmut*5**

(a) Confocal images of *N. benthamiana* leaves expressing full length SlSPCH-NeonGreen or the SlSPCHmut5 variant-NeonGreen driven by a 35Sx2 promoter. SlSPCH is primarily nuclear (black arrows). *SlSPCHmut5 is primarily nuclear at 22C,* but as temperature rises, fewer transformed cells (confirmed by expression of PM marker in grey) show nuclear localization of *SlSPCHmut5*. Scale bars represent 20µm. (b) Quantification of data depicted in (a), with fraction of transformed (PM-marker expressing) *N. benthamiana* epidermal cells that express SlSPCH variants in the nucleus as function of temperature (1.0 = 100%). Statistical tests in (b) represented as mean ± 95% confidence interval, each letter indicates a group of samples with no statistically significant difference between their means (Bonferroni corrected p value > 0.05). Sample sizes in (b) n = 3-7 0.4 $mm^2$ fields from 3 cotyledons.

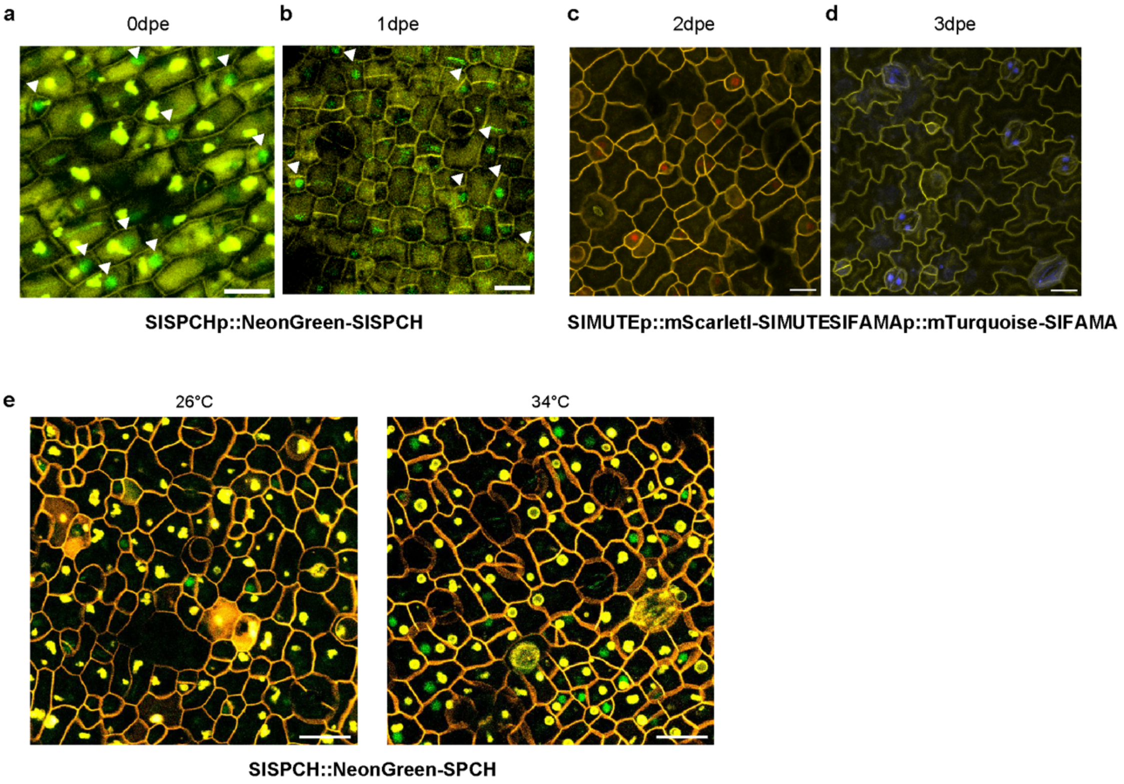


**Figure S4 Additional stomatal gene reporter expression**
(a-d) Confocal images of stomatal lineage reporters on days of peak expression in M82 cotyledons. Scale bars represent 20 µm. (e) Response of SlSPCHpro::NeonGreen-SlSPCH protein accumulation (green) at 0dpe in plants grown at either 26°C or 34°C. Cell outlines (yellow) are visualized with propidium iodide. Yellow inclusions in each cell are autofluorescence that is visible in multiple channels.

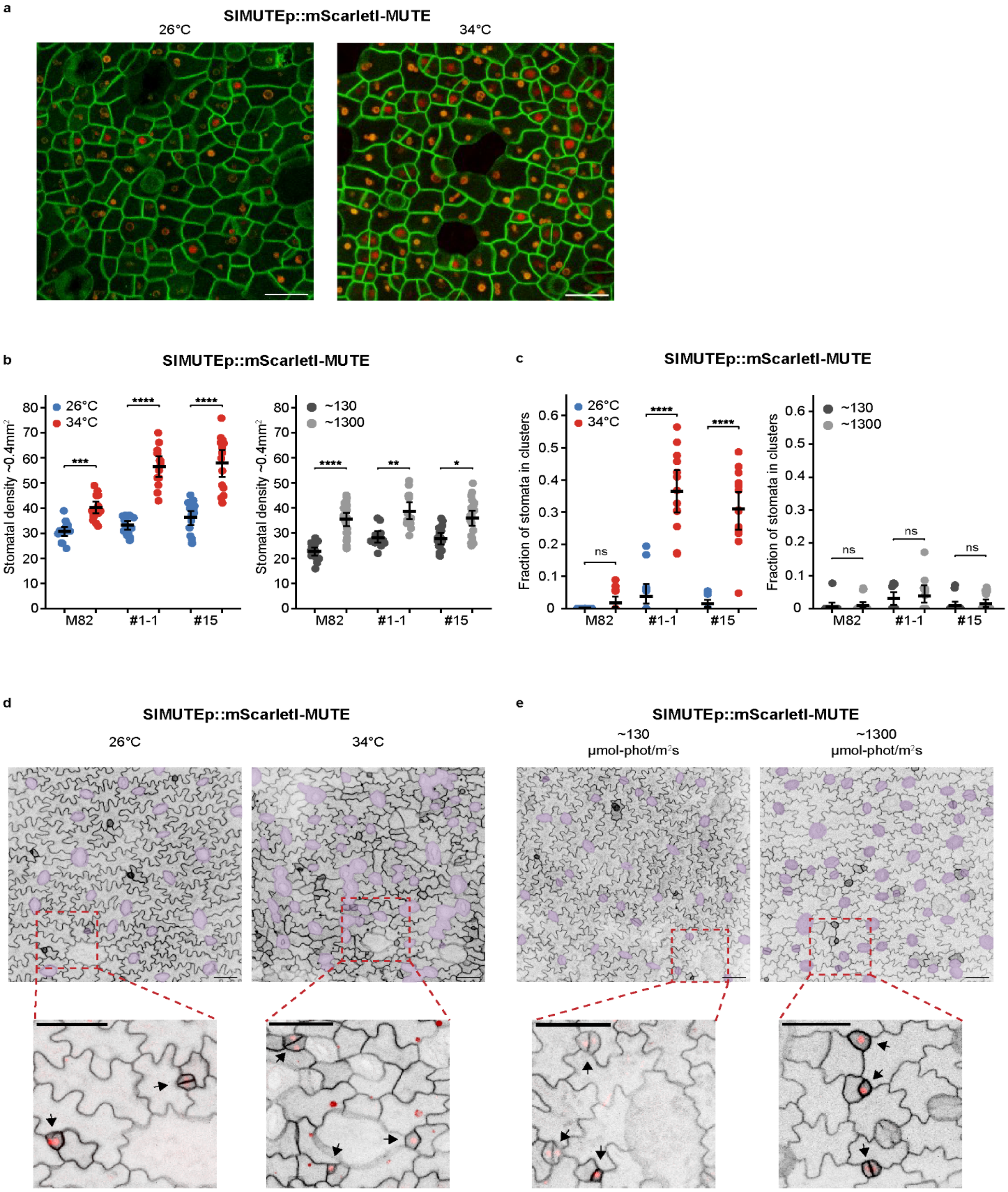


**Figure S5 Tomato lines bearing SlMUTE translational reporters display a hypersensitive response to increased temperature, but normal response to increased light**

(a) Confocal images of SlMUTEpro::mScarletI-MUTE (red) in 1dpe abaxial cotyledons grown at either 26°C or 34°C. Cell outlines (green) are visualized by propidium iodide. (b) Increase in stomatal density in two independent SlMUTEpro:mScarletI-SlMUTE (hereafter, SlMUTE) reporter lines (#1-1 and #15) in response to changes in temperature and light (c) Fraction of stomata found in clusters in SlMUTE, in response to changes in temperature and light. (d) Confocal images of 4-dpg cotyledons expressing SlMUTE at low and high light conditions. Stomata are false-colored purple. (e) Confocal images of 4-dpg cotyledons expressing SlMUTE at low and high temperature conditions. Stomata are false colored purple. Insets highlight clustered distribution of stomata consistent with "asymmetric divisions" incorrectly producing two stomata. Black arrows in insets point to cells expressing SlMUTE. Scale bars in (a) represent 30 µm. Scale bar in (e) and (f) represent 50 µm. Statistical tests in (b-c) are represented as mean ± 95% confidence interval. Bonferroni-corrected p values from Mann-Whitney U test are $**P < 0.01$; $****P < 0.0001$. n.s.: $P > 0.05$, not significant.  Sample sizes in (b-c) n = 5-15 0.4 $mm^2$ fields from 3-6 cotyledons.

### Table S2 Variation in SI of the tomato lines in response to changing environmental conditions

Table 2 shows the SI mean for each line and the inter-line statistics for each environmental condition. The statistical tests represented per condition as mean ± 95% confidence interval, each letter indicates a group of samples with no statistically significant difference between their means within environmental condition (Bonferroni corrected p value > 0.05). Sample sizes n = 8-20 0.4 mm2 fields from 4-8 cotyledons.

| Genotype | 100% RSWC | 30% RSWC | ~130 µmol-photons $m^{-2}$ $s^{-1}$ | ~1300 µmol-photons $m^{-2}$ $s^{-1}$ | 26°C | 34°C |
|---|---|---|---|---|---|---|
| M82 | 0.212 a | 0.188 a | 0.145 a | 0.183 a | 0.167 a | 0.177 a |
| spchp-cr#1 | 0.186 b | 0.167 b | 0.163 b | 0.185 a | 0.164 a | 0.174 a |
| spchp-cr#2 | 0.194 b | 0.183 ab | 0.149 ab | 0.176 a | 0.160 ab | 0.174 a |
| spchp-cr#3 | 0.184 b | 0.176 ab | 0.160 ab | 0.177 a | 0.167 a | 0.176 a |
| spchp-cr#4 | 0.178 b | 0.167 b | 0.125 c | 0.174 a | 0.152 b | 0.153 b |
| spchp-cr#5 | 0.165 c | 0.107 c | 0.117 c | 0. 125 b | 0.136 c | 0.035 c |

**Table S3 List of primers used for constructing reporters and for genotyping**

| Primer or gRNA ID | Sequence 5′-3′ | Purpose |
| --- | --- | --- |
| GGAG-SPCHp F | AAGAAGACTTGGAGTTCAACTTATAAAACTTTGGAATGGAGG | Golden Gate L0 promoter cloning |
| SPCHp-AATG R | AAGAAGACTTCATTTTTCAACGTTGAAAAAAGTAGAGGT | Golden Gate L0 promoter cloning |
| TTCG-SPCH F | AAGAAGACTTTTCGATGGATGGTGACCAAAATTTATCTG | Golden Gate L0 gene cloning |
| SPCH-mid F | TTGAAGACTTCCGTCTCTCCACGTATTCCTGGC | Golden Gate L0 gene cloning |
| SPCH-mid R | TTGAAGACTTACGGTtTTCAATATGACATTGGCGCC | Golden Gate L0 gene cloning |
| SPCH-GCTT R | TTGAAGACAAAAGCTTAGCAGAATGTCTGCTGAATCTGAT | Golden Gate L0 gene cloning |
| GGAG-MUTEp F | AAGAAGACTTGGAGTCTTAACCGTTTACAACTGAAAAGAATAAG | Golden Gate L0 promoter cloning |
| MUTEp-mid F | TTGAAGACTTAACGTcAGACCTATAATATATATTGTATATTGTCCAC | Golden Gate L0 promoter cloning |
| MUTEp-mid R | TTGAAGACTTCGTTTATTACCATATGGACATTTACGTT | Golden Gate L0 promoter cloning |
| MUTEp-AATG R | AAGAAGACTTCATTGATACTTTTTTTTTCTTCTTAAAAAAAAAAAACCTAAAT | Golden Gate L0 promoter cloning |
| TTCG-MUTE F | AAGAAGACTTTTCGATGTCTCACATAGCAGTGGAGAGAAACAGGAGgAGA | Golden Gate L0 gene cloning |
| MUTE-GCTT R | TTGAAGACAAAAGCCTATATCTCGTTGATACATAAAACATCAGATGAGGTGAA | Golden Gate L0 gene cloning |
| GGAG-FAMAp F | AAGAAGACTTGGAGATATTTGTTTCCTAGATATTTTTTTCCATCTCACA | Golden Gate L0 promoter cloning |

| | | |
|---|---|---|
| FAMAp-AATG R | AAGAAGACTTCATTTGTCTTGTTATAGTTTTTTTTTCTTTCTTTTTGTTTG | Golden Gate L0 promoter cloning |
| TTCG-FAMA F | AAGAAGACTTTTCGATGGAGAAAGAAGAAAATTGCCAGG | Golden Gate L0 gene cloning |
| FAMA-mid F | TTGAAGACTTTCATGCCTGGCTCCTATGTTC | Golden Gate L0 gene cloning |
| FAMA-mid R | TTGAAGACTTATGAGgGACCTCAATACACGAAGAT | Golden Gate L0 gene cloning |
| FAMA-C F | TTGAAGACTTGACCAGGACAACTCATAAAGGC | Golden Gate L0 gene cloning |
| FAMA-C R | TTGAAGACTTGGTCTcCTTCTTGATAGAATCTTGATCAT | Golden Gate L0 gene cloning |
| FAMA-GCTT R | TTGAAGACAAAAGCCTAGTTGATATCATATGGGGCTATTTCAGC | Golden Gate L0 gene cloning |

**Table S4 List of sgRNAs used for mutagenesis and PCR primers used for genotyping**

| Primer or gRNA ID | Sequence 5′-3′ | Purpose |
|---|---|---|
| gRNA1 | TTCAACGTTGAAAAAAGTAG | Golden Gate cloning |
| gRNA2 | TCTGCACTTTTACTTTTGTT | Golden Gate cloning |
| gRNA3 | ATATATGCCATTTTGTTGAG | Golden Gate cloning |
| gRNA4 | TTATGTTAGCAAGTTCAAAC | Golden Gate cloning |
| gRNA5 | TTTCAAATAGAAAGATACGA | Golden Gate cloning |
| gRNA6 | ATATAATCTACACATTAACA | Golden Gate cloning |
| gRNA7 | ATACTCATACTTCATAGTTA | Golden Gate cloning |
| gRNA8 | GTTTGGCTCTGCTTGAAAGC | Golden Gate cloning |
| gRNA9 | ACTTAAAAAGAAACGCGACG | Golden Gate cloning |
| gRNA10 | ATAAATATGATATATTATGG | Golden Gate cloning |
| gRNA11 | GGGAATAAGAGAGAGAGCAA | Golden Gate cloning |
| SlSPCHpro F | TCTCGTTTACGGAAAGTAGAAACC | Genotyping, cloning and sequencing |
| SPCH155 R | ATTGTTGCAGTCTGGTTGTTGAT | Genotyping, cloning and sequencing |
| SlSPCH2937up F | GCGTCGTCGGAAAATCCAAG | Promoter sequencing |
| SlSPCH2168up F | GAGCTTGTTTGGCTCTGCTTG | Promoter sequencing |
| SlSPCH1700up F | TGCTACATTAATTACATTGAGTGAGAA | Promoter sequencing |
| SlSPCH1311up F | CAAATAGAAAGATACGAAGGTCGAAGA | Promoter sequencing |
| SlSPCH943up F | AAGAACCCGTTTGAACTTGC | Promoter sequencing |
| SlSPCH F | TATTCCCTTTTCTACATCTCCTCTTT | Genotyping, cloning and sequencing |

| | | |
|---|---|---|
| SlSPCH_R | TTCTCTTATTATTGTTACAGCTCCCTA | Genotyping, cloning and sequencing |
| SlSPCH200 | GGTGTCAGAAGAAGGAAAGAAGA | Gene sequencing |
| SlSPCH386 | ACTTTGATGCCTTGTTTTTATGCTAA | Gene sequencing |
| SlSPCH951 | GGTGATCAAGCATCAATAATTGGT | Gene sequencing |
| SPCH1371 | CCGTCTCTCCACGTATTCCTGGC | Gene sequencing |
